# A 28-color panel for classical and non-classical T lymphocytes in decidua and PBMC in rhesus macaques

**DOI:** 10.64898/2026.08.27.747578

**Authors:** Matilda J. Moström, Amitinder Kaur, Marissa Fahlberg

## Abstract

**Panel Presentation:** *Purpose and appropriate sample types:* This 28-color panel was developed to identify classical and non-classical T lymphocytes in decidual leukocytes and peripheral blood mononuclear cells (PBMC) of pregnant rhesus macaques (**Table 1**). By profiling these T lymphocytes, we can investigate how maternal immunity balances tolerance to fetal antigens with protection against vertically transmitted pathogens. The selected markers define memory populations and characterize tissue residency, activation, proliferation, cytotoxicity, trafficking, and exhaustion status. This panel also delineates B lymphocytes and NK cells to confirm expected frequencies. The utility of this panel is aimed at evaluating cellular immune correlates of protection against congenital infections at the maternal-fetal interface and PBMC in rhesus macaques.

## Background

The maternal-fetal interface is a unique environment composed of the placenta (fetal-origin) and the decidua (maternal-origin) that serves several essential functions during pregnancy. These functions include nutrient and gas exchange between the maternal and fetal circulations, immune tolerance to paternal antigens in the fetus and placenta, and protection of the fetus against maternal pathogens. To date, understanding of maternal-fetal immunology in humans has focused on reproductive research, whereas data on immune perturbations during viral congenital infections are sparse. Unlike in reproductive research where samples can be obtained during delivery, acquiring human samples to understand congenital pathogen transmission is challenging since placental studies are not routine and mothers with infections are often undiagnosed. Thus, using a closely related animal model such as rhesus macaques to characterize the immunological environment of the maternal-fetal interface during maternal infection is a powerful approach to improving our knowledge of protective immune correlates of congenital transmission and fetal infection.

We recently reported on immune cell characterization of the decidual maternal-fetal interface in normal and Zika virus-infected rhesus macaque dams using 28-color and 18-color flow cytometry panels (1). In this OMIP, we describe development, optimization, and further improvement of the 28-color flow cytometry panel optimized to phenotype classical and non-classical T lymphocytes in cryopreserved decidual leukocytes. It resolves T cell memory, activation, exhaustion, cytotoxicity, and proliferation potential of T lymphocytes. This panel also includes limited B and NK cell phenotyping as an internal control to confirm expected immune cell distributions and assess peripheral blood contamination. In rhesus macaques, B cells and NK cells respectively comprise approximately 10-20% and 5-15% of CD45^+^ leukocytes in blood, whereas in decidua these frequencies are typically <5% and >50% indicating tissue specificity and minimal blood contamination during processing and analysis (1, 2).

We designed this panel to analyze T lymphocytes and enable investigation of tissue-level cellular immunity probing pathogen- and vaccine-inducible responses (**Figures 1-2**, **Table 2**). As naïve T lymphocytes encounter their cognate antigen in secondary lymphoid organs, they undergo differentiation into varying types of memory subsets (3) and travel into tissues that include the decidua (4–7). In this panel, the markers CD45RA, CD95, CD28, and CCR5 are used to define five memory subsets in rhesus macaques. These include naïve (CD95^-^ CD28^+^ CCR5^-^ CD45RA^+^), central memory (CM; CD95^+^ CD28^+^ CCR5^-^ CD45RA^-^), transitional effector memory (TEM; CD95^+^ CD28^+^ CCR5^+^ CD45RA^-^), effector memory (EM; CD95^+^ CD28^-^ CCR5^+/–^ CD45RA^-^), and terminally differentiated effector memory (TEMRA; CD95^+^ CD28^-^ CCR5^+/–^ CD45RA^+^) (1, 8–10). CD4^+^ T lymphocytes can be further classified into T helper (Th) subsets, including Th1, Th2, and Th17, that respond uniquely to intracellular and extracellular pathogens of different types (viruses, bacteria, fungi, and parasites) (11, 12). These T helper cell subtypes can be defined by their chemokine receptor expression patterns or cytokine secretion profile (13). For this purpose, antibodies binding to the chemokine receptors CCR4, CCR5, CCR6, and CXCR3 were included in this panel. Among CD4^+^ T lymphocytes, CXCR3^+^ CCR6^-^ CCR4^-^ are associated with a Th1 phenotype, CXCR3^-^ CCR6^-^ CCR4^+^ are associated with a Th2 phenotype, and CXCR3^-^ CCR6^+^ CCR4^+^ are associated with a Th17 phenotype (13, 14). In addition to T helper-defining chemokine receptors, we also included the chemokine receptor CX3CR1, associated with T cell chemotaxis to sites of endothelial inflammation (15, 16).

**Figure 1.**
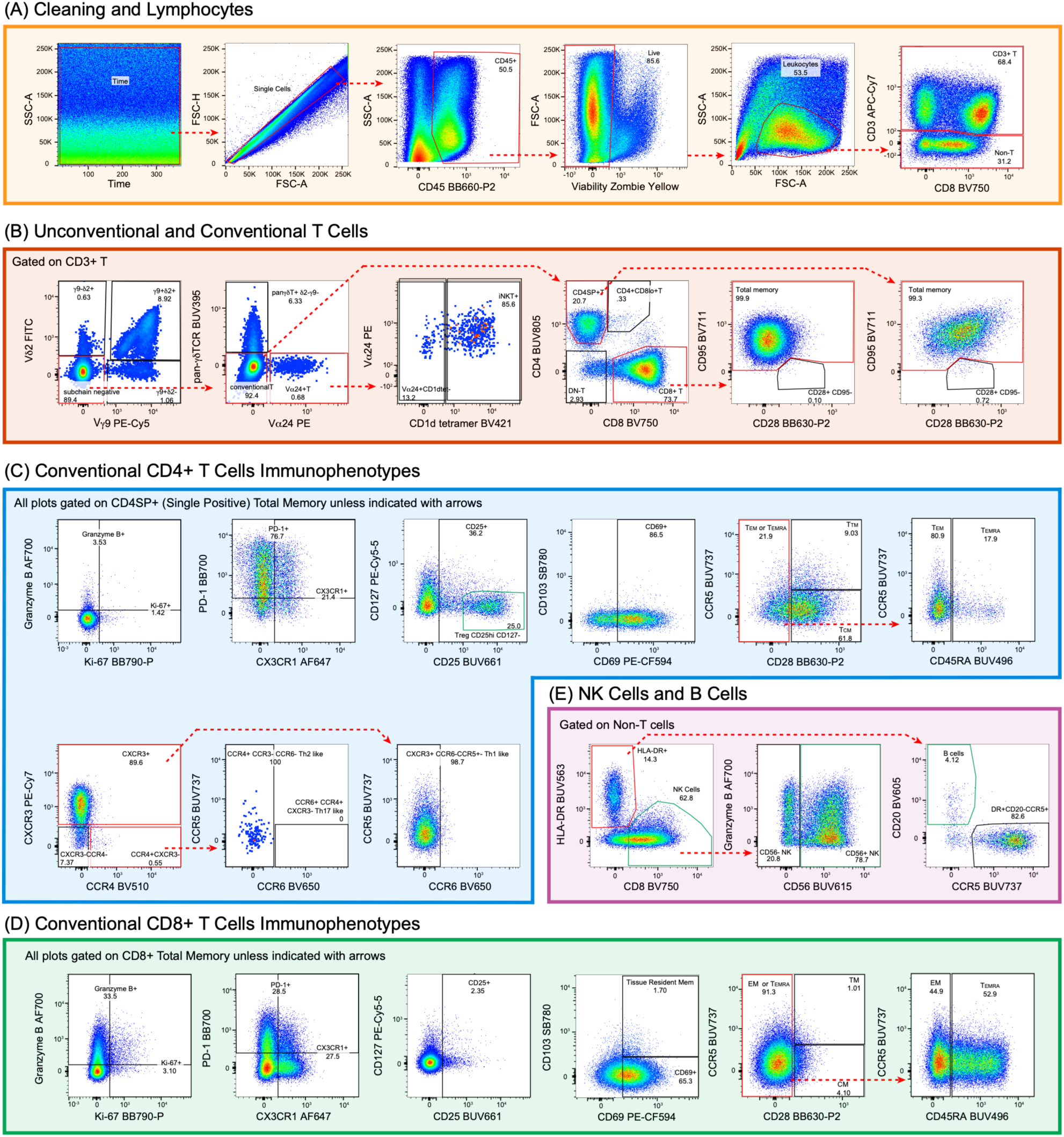
Example gating strategy of decidual leukocytes. A) Samples are gated on Time vs SSC-A to remove fluidic disturbances including air contamination and micro-clogs. Next, doublets are removed and live CD45^+^ cells are gated followed by a leukocyte gate based on size and complexity characteristics (FSC-A vs SSC-A). The leukocytes gate is used to subset CD3^+^ and CD3^-^ lymphocytes for further differentiation in boxes B and E. B) CD3^+^ T lymphocytes are gated to classify non-classical and classical T lymphocytes including γδ T, iNKT, CD4SP^+^ (<u>S</u>ingle <u>P</u>ositive), CD4+CD8lo+ T, and CD8^+^ T lymphocytes. C) Conventional CD4SP^+^ total memory lymphocytes are gated to characterize expression of activation, exhaustion, inflammation, trafficking, and whether they are Th1-like, Th2-like, or Th17-like. D) Conventional CD8^+^ total memory lymphocytes are gated to characterize expression of markers as in CD4 excluding the helper subsets. E) NK and B cell gates are used to confirm expected cell frequencies in the decidua. Note: gating positions are placed based on biological comparison to PBMC sample (Figure 2) and FMO controls (Figure 13). Red gates denote “parent” gates that have corresponding hierarchical child gates, while black gates are terminal child gates. Green gates highlight cell populations used as internal controls to confirm expected cell frequencies due to known biological differences between PBMC and decidua. These include CD25hi CD4^+^ T lymphocytes, B cells, NK cells, and CD56 expression on NK cells.

**Figure 2.**
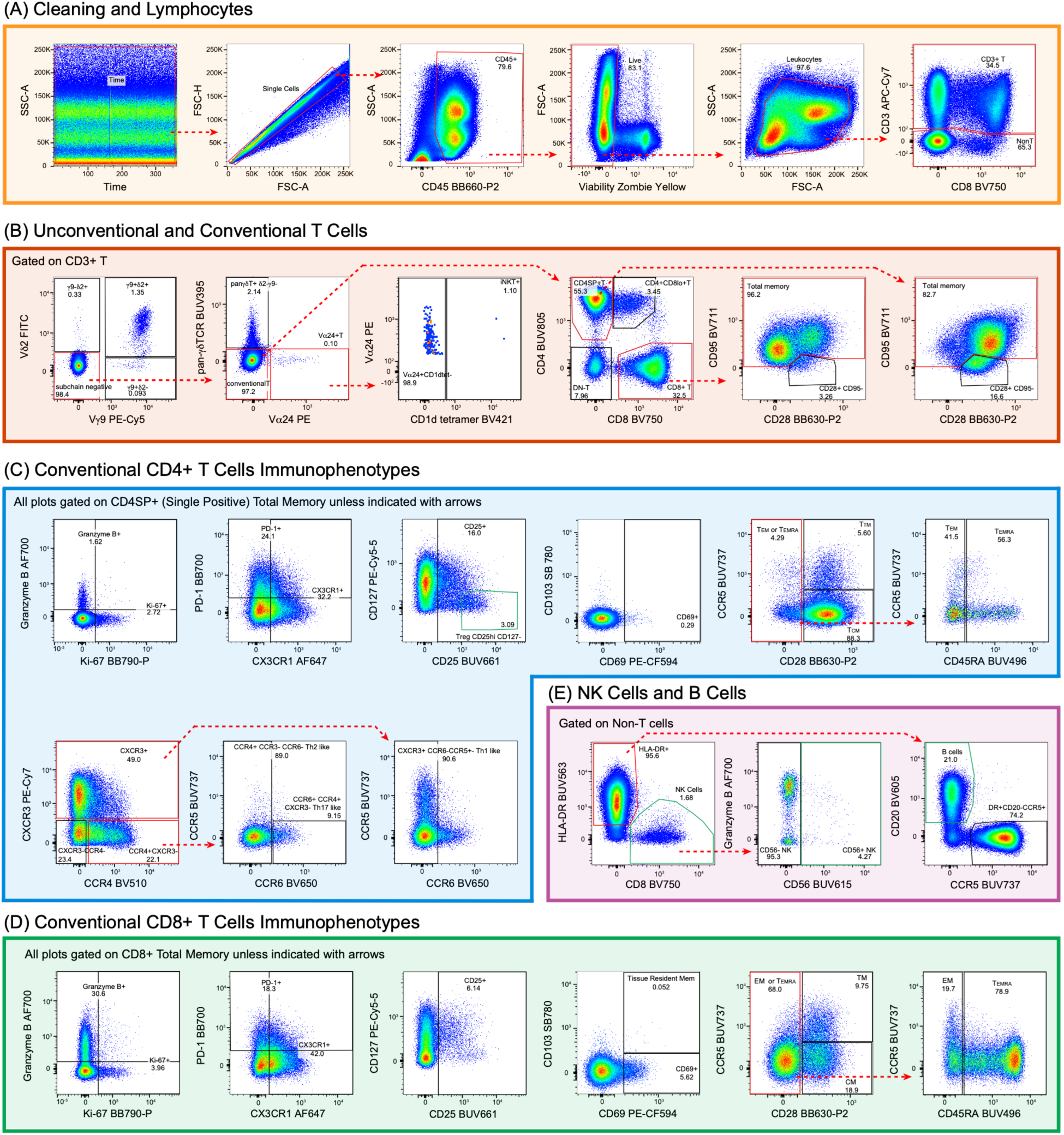
PBMC gating hierarchy. The gating strategy used for decidual leukocytes in Figure 1 is applied with minor sample-type adjustments to a representative PBMC sample from a pregnant rhesus macaque at similar gestation.

**Table 1.** Summary table for application of OMIP-XXX.

|  |  |
| --- | --- |
| Purpose | Phenotype and function of adaptive and innate T lymphocytes in the decidua and PBMCs |
| Species | Indian-origin rhesus macaque |
| Cell type | Decidual leukocytes and PBMC |
| Cross references | OMIP-005, OMIP-052, OMIP-075 |

**Table 2.**
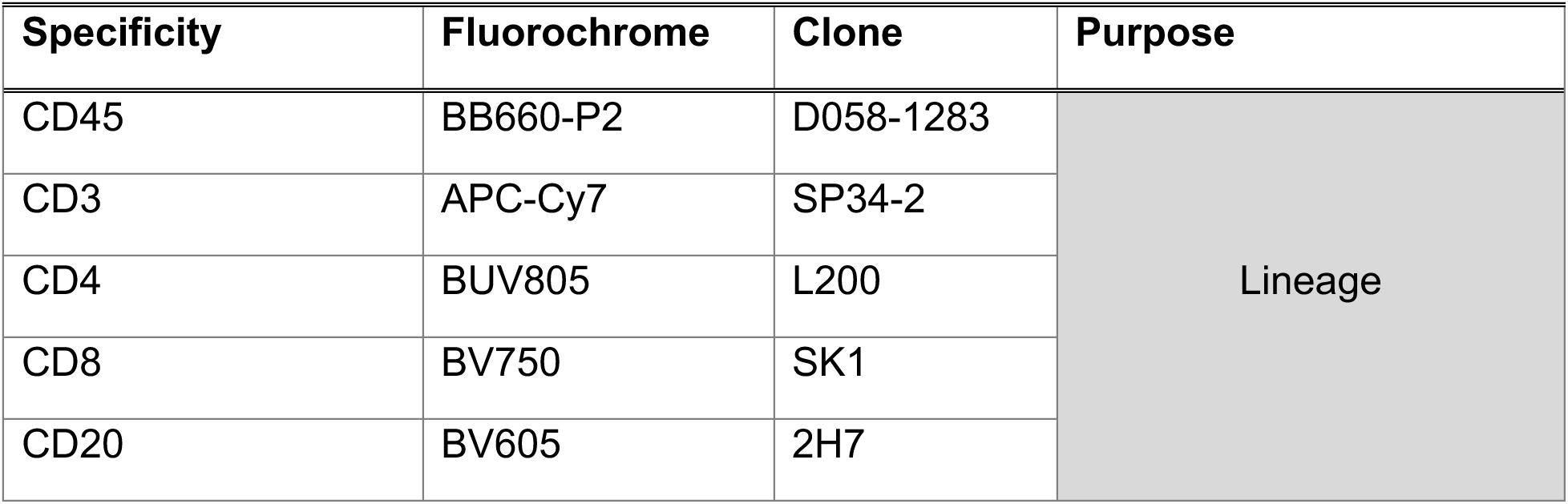
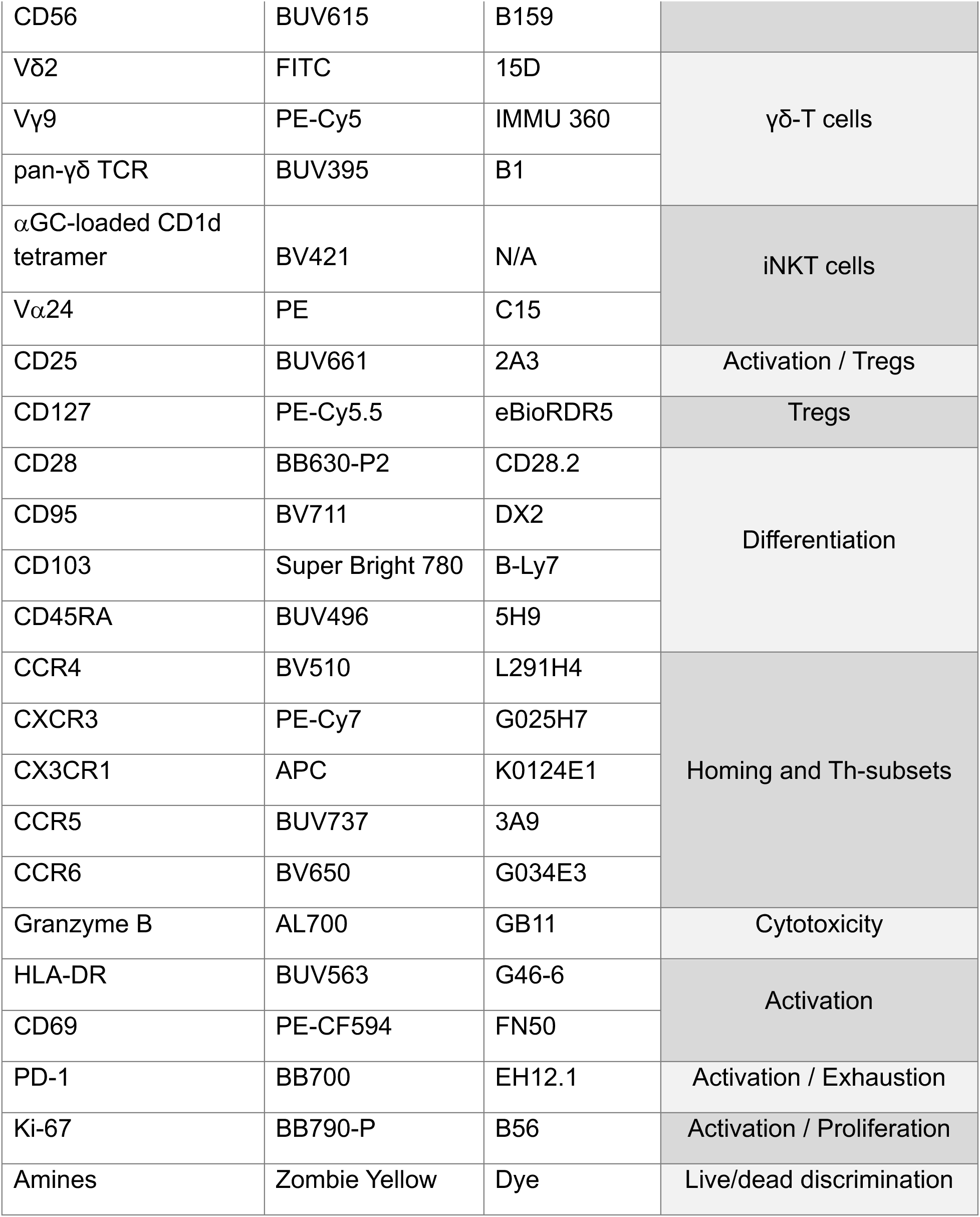
Reagents used for OMIP-XXX.

An additional CD4^+^ T cell type that plays an important role in infection is the T regulatory cell (Treg) which modulates the magnitude of immune responses (17, 18). In the decidua, Tregs play a central role in mediating tolerance in healthy pregnancies, and their disruption is associated with several adverse pregnancy outcomes, including recurring spontaneous abortions and pre-eclampsia (19–23). We define Tregs in this panel as CD4^+^ CD25^hi^ and CD127^low^, which has been validated in rhesus macaques to correspond to the same CD4^+^ CD25^hi^ lymphocytes that co-express the nuclear transcription factor FOXP3, a canonical marker of Tregs (24).

Beyond the classical phenotypes described, some T lymphocytes are “non-classical” and mount rapid and broad innate-like immune responses (25–28). In this panel, we included markers that identify two unconventional “innate” T lymphocyte subsets: invariant natural killer T (iNKT) and gamma delta (γδ) T lymphocytes. iNKTs have a semi-invariant TCR that recognizes glycolipid antigens presented on the nonpolymorphic MHC class I-like CD1d molecule, and γδ T cells recognize a variety of antigen types in receptor-dependent or -independent mechanisms (26–28). Both of these non-classical T lymphocyte subsets are present at the maternal-fetal interface and may regulate pregnancy. For example, in early gestation, decidual γδ T lymphocytes differ in frequency and TCR repertoire compared to peripheral blood (29, 30). iNKTs are identified in non-human primates as CD3^+^ T lymphocytes that express the Vα24 TCR (Vα24+) and bind to CD1d tetramers loaded with α-galactosyl-ceramide (αGC) or its analog PBS-57 (31, 32). To identify γδ T cells in non-human primates, we included two antibodies targeting the variable δ2 (Vδ2) and γ9 chains (Vγ9), and one pan-γδ TCR antibody. Notably, the double-positive Vδ2+ and Vγ9+ T cell subsets were pan-γδTCR low and would likely be missed without the chain-specific antibodies. However, we still included the pan-γδ TCR antibody to capture any additional γδ T cells which did express this marker without a Vγ9 or Vδ2 chain.

Last, we included several surface markers associated with cellular states and functions. Chemokine receptors CXCR3, CCR4, CCR5, CCR6, and CX3CR1 are markers of lymphocyte trafficking (33, 34), Granzyme B is a marker of cytolytic potential (35–37), PD-1 is an activation and exhaustion marker (38, 39), CD69, CD25, and HLA-DR are activation markers (40–43), and CD103 and CD69 define tissue residency (44). The choice of these markers was based on phenotypes observed in inflammation and infection as well as reagent availability. Some markers that are relevant to T lymphocytes in humans, such as CD57, were excluded because they lack cross-reactive antibody clones in rhesus macaques.

Successful implementation of this panel required several practical considerations, most notably the addition of antibodies and antibody cocktails across six sequential staining steps. Each antibody cocktail was prepared either the same day or the previous day, in an appropriate cocktail buffer for the reagents. The six antibody addition steps consisted of: (1) viability dye staining, (2) iNKT markers, (3–4) chemokine and cytokine receptor staining, (5) surface marker staining, and (6) intracellular marker staining. Chemokine and cytokine receptor staining was performed at 37°C, whereas all other staining steps were conducted at room temperature. In addition, intracellular marker detection required fixation and permeabilization. Details of the staining procedure can be found in the protocol and **Figure 3**. Collectively, the total thawing, staining, fixation, and permeabilization steps resulted in a total processing time of approximately 9-12 hours.

**Figure 3.**
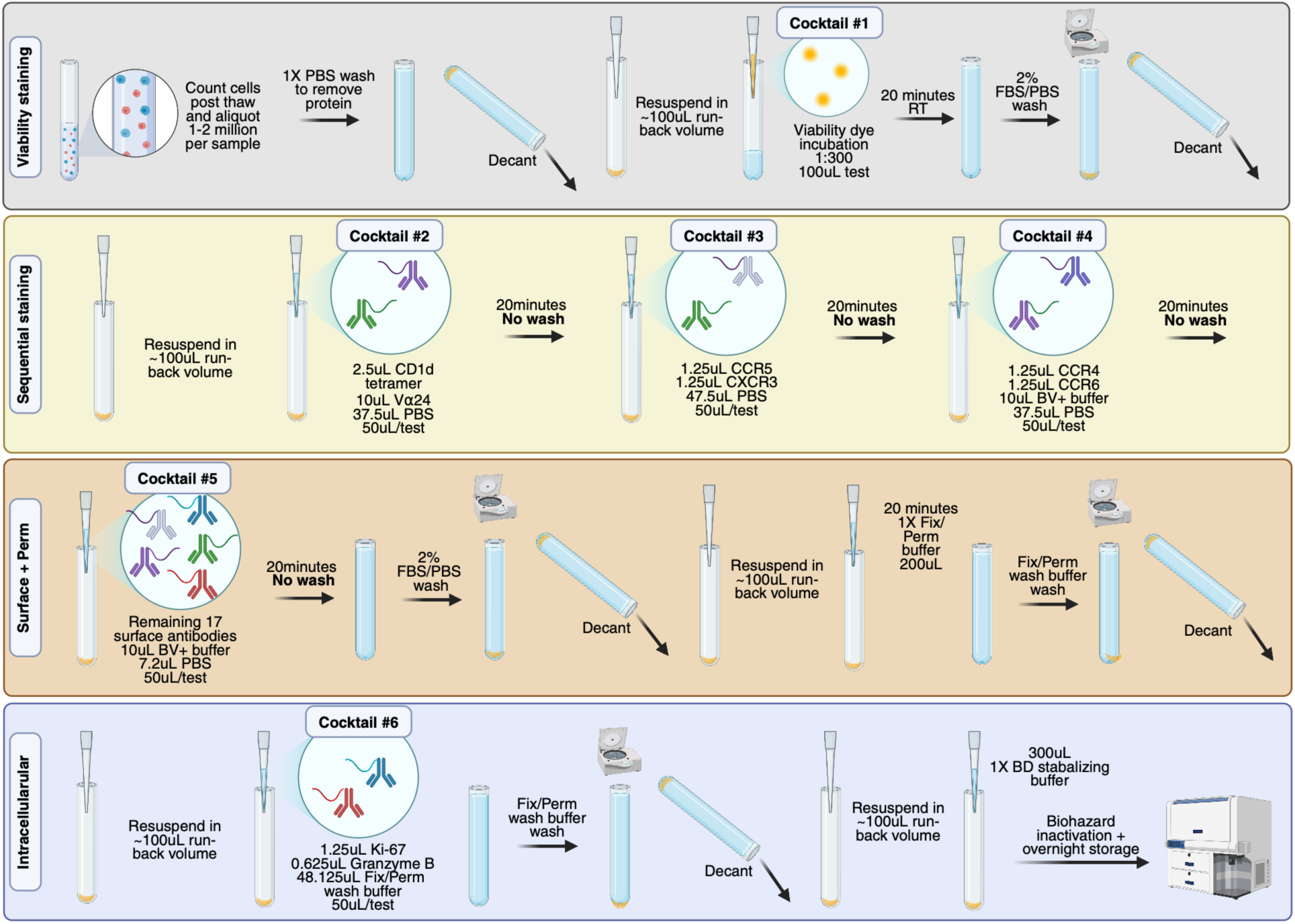
Visual description of staining procedure. The full procedure for staining this panel is shown. Antibody cocktail numbers correspond to those in **Table 4**. Centrifugation settings, buffer recipes, and additional details can be viewed in the step-by-step protocol.

Due to the length of the sequential staining protocol, it was not feasible to retrieve cryopreserved samples from liquid nitrogen storage, thaw cells, prepare antibody cocktails, complete staining, and acquire samples on the same day. Therefore, the panel was optimized incorporating several practical workflow decisions, including: (1) preparation of several antibody cocktails one day prior to staining, (2) transfer of cryovials from liquid nitrogen storage to a −80°C freezer one day prior to thawing, and (3) next-day sample acquisition following completion of staining.

Antibody cocktail storage was evaluated directly and demonstrated comparable staining performance after one day of storage relative to freshly prepared cocktails (**Figure 4**), whereas reduced staining quality was observed after three days of storage (data not shown). Based on these findings, several antibody cocktails were prepared one day prior to staining. This strategy is similar to the workflow described by Liechti et al. which enabled de-risking precious samples using a quality control stain of prepared cocktails (45). Likewise, robust panel performance was observed following the intermediate −80°C storage step prior to thawing. While this handling procedure may result in modest differences compared to direct thawing from liquid nitrogen, any effects are expected to be comparable to those associated with routine transport of cryopreserved samples on dry ice before or after processing. Finally, although sample acquisition was performed on the day following completion of staining, samples were in practice acquired within 12-16 hours of stain completion. Longer post-stain storage intervals may require additional optimization.

**Figure 4.**
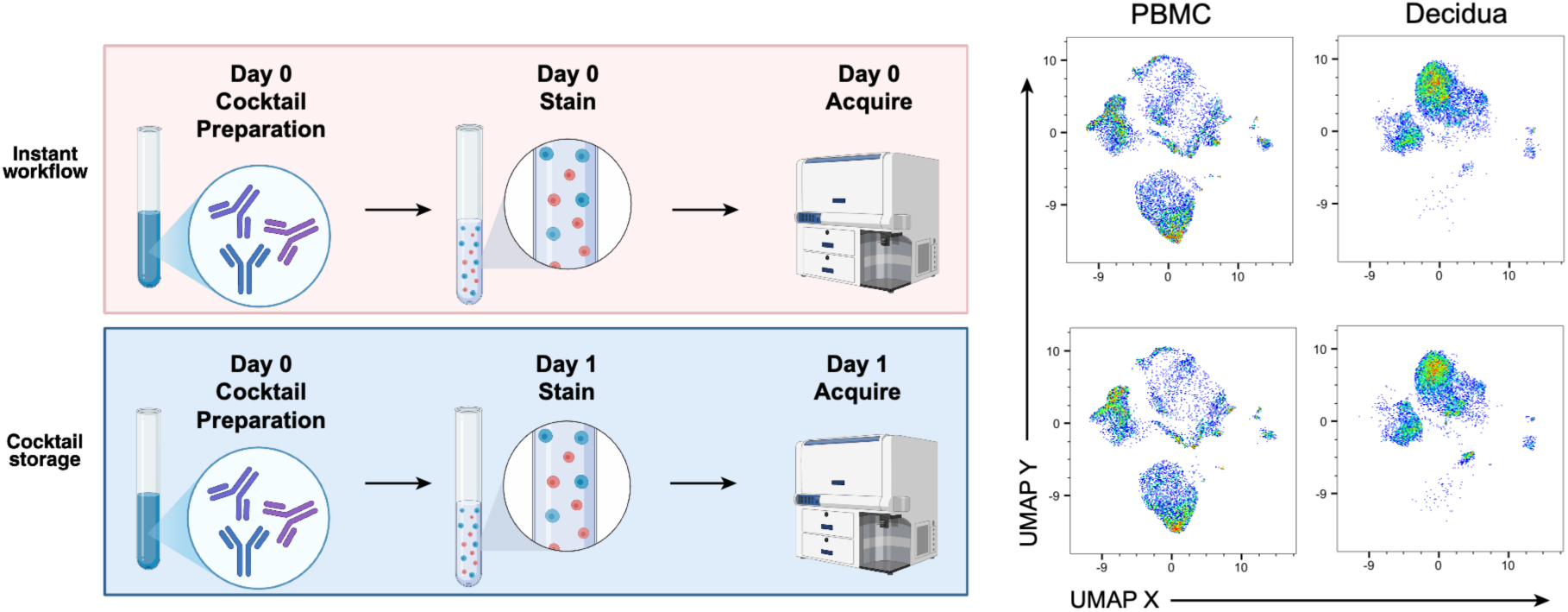
Impact of one-day antibody cocktail storage on cell population resolution. The schematic represents the workflow of an experiment comparing one-day antibody cocktail storage to same-day cocktailing, staining, and acquisition using PBMC and decidua from two animals (one animal per column). The conditions tested were: 1) preparation of antibody cocktail, antibody staining, and sample acquisition all on the same day (top row, instant workflow); and 2) preparation of antibody cocktail one day prior to antibody staining and sample acquisition (bottom row, cocktail storage workflow). On the right, UMAPs of 10,000 live single CD45+ CD3+ or CD20+ cells depict global cellular structure. UMAPs were generated with all fluorescent parameters except those used for gating, concatenating samples, and using default UMAP settings in FlowJo.

This panel was acquired on the BD FACSymphony A5 (**Table 3**), and bead-based particles were found to be suitable for single stain control to calculate compensation. However, the use of bright beads was imperative to accurately compensate samples, as many of the markers stained brightly on samples and insufficiently bright compensation beads resulted in false positives and high background, described further in the Panel Design and Optimization strategy.

**Table 3.** Instrument configuration for BD FACSymphony A5.

| Laser wavelength [nm] | Laser power [mW] | Laser type | Spectral range for detector [nm] | Optical filters |  |  | Fluorochrome |
| --- | --- | --- | --- | --- | --- | --- | --- |
|  |  |  |  | Dichroic #1 [nm] | Dichroic #2 [nm] | Band pass [nm] |  |
| 488 | 200 | DPSS | 750-810 | - | 750 | 780/60 | BB790-P |
|  |  |  | 685-735 | 750 | 690 | 710/50 | BB700 |
|  |  |  | 655-685 | 690 | 635 | 670/30 | BB660-P2 |
|  |  |  | 600-620 | 635 | 600 | 610/20 | BB630-P2 |
|  |  |  | 562.5-587.5 | 600 | 500 | 515/20 | FITC |
| 405 | 200 | DPSS | 750-810 | - | 750 | 780/60 | SuperBright 780 |
|  |  |  | 722.5-757.5 | 750 | 710 | 740/35 | BV750 |
|  |  |  | 685-735 | 710 | 690 | 710/50 | BV711 |
|  |  |  | 655-685 | 690 | 635 | 670/30 | BV650 |
|  |  |  | 600-620 | 635 | 600 | 610/20 | BV605 |
|  |  |  | 577.5-592.5 | 600 | 550 | 585/15 | Zombie Yellow |
|  |  |  | 500-550 | 550 | 505 | 525/50 | BV510 |
|  |  |  | 425-475 | 505 | 410 | 450/50 | BV421 |
| 355 | 60 | DPSS | 790-850 | - | 770 | 820/60 | BUV805 |
|  |  |  | 722.5-757.5 | 770 | 690 | 740/35 | BUV737 |
|  |  |  | 655-685 | 690 | 635 | 670/30 | BUV661 |
|  |  |  | 600-620 | 635 | 600 | 610/20 | BUV615 |
|  |  |  | 578.5-593.5 | 600 | 550 | 586/15 | BUV563 |
|  |  |  | 500-530 | 550 | 450 | 515/30 | BUV496 |
|  |  |  | 365-393 | 450 | - | 379/28 | BUV395 |
| 561 | 200 | DPSS | 750-810 | - | 750 | 780/60 | PE-Cy7 |
|  |  |  | 685-735 | 750 | 690 | 710/50 | PE-Cy5.5 |
|  |  |  | 655-685 | 690 | 635 | 670/30 | PE-Cy5 |
|  |  |  | 600-620 | 635 | 600 | 610/20 | PE-CF594 |
|  |  |  | 578.5-593.5 | 600 | 580 | 586/15 | PE |
| 628 | 200 | DPSS | 750-810 | - | 750 | 780/60 | APC-Cy7 |
|  |  |  | 685-735 | 750 | 690 | 710/50 | AF700 |
|  |  |  | 655-685 | 690 | 665 | 670/30 | AF647 |

A conservative gating strategy was used to define the populations described herein, although gates could be adjusted for continuous populations depending on the context of use. Several data-cleaning steps were first applied to remove artifacts, dead cells, and doublets. These included exclusion of fluidic disturbances with a time gate, followed by sequential singlet gating using FSC-H vs FSC-A and SSA-A vs SSC-H. Viable CD45+ white blood cells were then identified, followed by leukocyte gating and separation of T lymphocytes vs non-T lymphocytes using CD3 and CD8 expression. Gate thresholds were established using a combination of marker expression patterns in PBMC and decidua, expected biological distributions, and FMO controls. To improve analysis reproducibility, gates extended to the axis limits to ensure inclusion of all qualifying events (46).

Notably, two reagents used in this panel and provided by BD Biosciences were discontinued since initial panel development (CD28 clone CD28.2 BB630-P2 and CD45 clone D058-1283 BB660-P2). These reagents have been updated by Waters Biosciences with new fluorochromes (RB613 and RB670, respectively). Implementation of this panel with updated fluorochromes would require a panel re-evaluation for equivalent or superior performance with spectrally cleaner dyes.

### Similarities to other OMIPs

Panels related to T lymphocyte characterization in rhesus macaques are found in OMIP-005 and OMIP-052. Both investigate cytokine staining following antigen stimulation in PBMC and do not include broad immunophenotyping markers. OMIP-075, developed for humans and non-human primates, focuses on both intracellular cytokine staining and broad T lymphocyte immunophenotyping markers for PBMC. However, this OMIP is optimized for cynomolgus macaques and not rhesus, and clone cross-reactivity can vary greatly by species. This is the first decidual OMIP for any species and the first in-depth broad T lymphocyte immunophenotyping panel in rhesus macaques.

### Technical Information

All samples were acquired on the BD FACSymphony A5, with the configuration shown in **Table 3**. Reagents used in the final panel are described in **Table 4**, and a detailed protocol is written for reproduction of this panel.

**Table 4.** Reagent information used in OMIP-XXX. This table provides a detailed description of the fluorescent reagents used in the panel. Antibody cocktails are prepared based on a 50μL per test volume, except for the viability dye* which is prepared as a 100μL test volume. The antibody cocktails are added to each sample after decanting and/or without washing in the indicated sequence and are visually contextualized in **Figure 11**. **Catalog number represents a custom conjugate available from BD Biosciences High Parameter Solutions. “Intra” stands for intracellular stain.

| Marker | Fluorochrome | Clone | Catalog# | Lot Number | Manufacturer | Antibody cocktail number | µL antibody (per 50µL/test cocktail) | Titer (per 50µL/test cocktail) | RRID |
| --- | --- | --- | --- | --- | --- | --- | --- | --- | --- |
| Amines | Zombie Yellow | Dye | 423104 | NA | Biolegend | 1 | *0.333 | 1:300 | NA |
| αGC-loaded CD1d tetramer | BV421 | Tetramer | NA | NA | NIH tetramer core | 2 | 10 | 1:5 | Tetramer core SCR_026557 |
| Vα24 | PE | C15 | IM2283 | 200060 | Beckman-Coulter | 2 | 2.5 | 1:40 | AB_131321 |
| CCR5 | BUV737 | 3A9 | 748873 | 1034541 | BD Biosciences | 3 | 1.25 | 1:40 | AB_2873276 |
| CXCR3 | PE-Cy7 | G025H7 | 353720 | B300691 | Biolegend | 3 | 1.25 | 1:40 | AB_11219383 |
| CCR4 | BV510 | L291H4 | 359415 | B325088 | Biolegend | 4 | 1.25 | 1:40 | AB_2562436 |
| CCR6 | BV650 | G034E3 | 353426 | B307432 | Biolegend | 4 | 1.25 | 1:40 | AB_2563869 |
| CD103 | Super Bright 780 | B-Ly7 | 78-1038-41 | 2228658 | Invitrogen | 5 | 5 | 1:10 | AB_2762587 |
| CD127 | PE-Cy5.5 | eBioRDR5 | 35-1278-42 | 2298663 | Invitrogen | 5 | 5 | 1:10 | AB_2744722 |
| CD20 | BV605 | 2H7 | 302334 | B243544 | Biolegend | 5 | 0.625 | 1:80 | AB_2563398 |
| CD25 | BUV661 | 2A3 | 741685 | 1026937 | BD Biosciences | 5 | 1.25 | 1:40 | AB_2871068 |
| CD28 | BB630-P2 | CD28.2 | **624294 | 1013555 | BD Biosciences | 5 | 1.25 | 1:40 | AB_3751209 |
| CD3 | APC-Cy7 | SP34-2 | 557757 | 223215 | BD Biosciences | 5 | 2.5 | 1:20 | AB_396863 |
| CD4 | BUV805 | L200 | 749212 | 1022167 | BD Biosciences | 5 | 0.625 | 1:80 | AB_2873590 |
| CD45 | BB660-P2 | D058-1283 | **624295 | 1029398 | BD Biosciences | 5 | 0.15 | 1:320 | AB_3751210 |
| CD45RA | BUV496 | 5H9 | 741182 | 1026942 | BD Biosciences | 5 | 0.625 | 1:80 | AB_2870749 |
| CD56 | BUV615 | B159 | 751349 | 1026934 | BD Biosciences | 5 | 2.5 | 1:20 | AB_2875357 |
| CD69 | PE-CF594 | FN50 | 562645 | 9030547 | BD Biosciences | 5 | 1.25 | 1:40 | AB_2737699 |
| CD8 | BV750 | SK1 | 344756 | B300774 | Biolegend | 5 | 2.5 | 1:20 | AB_2810547 |
| CD95 | BV711 | DX2 | 305644 | B300083 | Biolegend | 5 | 1.25 | 1:40 | AB_2632623 |
| CX3CR1 | APC | K012E1 | Custom | B331099 | Biolegend | 5 | 5 | 1:10 | AB_3751213 |
| HLA-DR | BUV563 | G46-6 | 748340 | 1013153 | BD Biosciences | 5 | 0.625 | 1:160 | AB_2872759 |
| pan-γδTCR | BUV395 | B1 | 564155 | 325885 | BD Biosciences | 5 | 5 | 1:10 | AB_2738627 |
| PD-1 | BB700 | EH12.1 | 566461 | 344583 | BD Biosciences | 5 | 0.625 | 1:80 | AB_3751212 |
| Vγ9 | PE-Cy5 | IMMU 360 | A6363 | 200029 | Beckman-Coulter | 5 | 0.625 | 1:80 | AB_3644200 |
| Vδ2 | FITC | 15D | Custom | NA | Creative Biolabs | 5 | 2 | 1:25 | AB_3751211 |
| Granzyme B | AF700 | GB11 | 560213 | 195628 | BD Biosciences | 6 (Intra) | 0.625 | 1:80 | AB_1645453 |
| Ki-67 | BB790-P | B56 | **624296 | 1014847 | BD Biosciences | 6 (Intra) | 1.25 | 1:40 | AB_3751214 |

## Materials

- 1X Phosphate Buffered Saline (PBS) (Corning)
- RPMI 1640 with phenol red (Gibco)
- Penicillin 100X (Corning)
- Streptomycin 100X (Corning)
- Heat-inactivated Fetal Bovine Serum (FBS) (Gibco)
- HEPES 1M (Gibco)
- L-Glutamine 200mM (Sigma)
- DNase I from bovine pancreas (Millipore Sigma)
- ViaStain AOPI Staining Solution (Nexcelom Bioscience/Revvity)
- Brilliant Stain Buffer Plus (BD Biosciences, RRID: 2869761)
- Stabilizing Fixative 3X Concentrate; diluted to 1X with distilled water (BD Biosciences)
- BD Cytofix/Cytoperm™ Fixation/Permeabilization Kit (BD Biosciences)
- ArC Amine Reactive Compensation Bead Kit (Invitrogen)
- Compensation beads (BioLegend)
- Surface and intracellular antibody cocktails (see **Table 4**)
- Zombie Yellow™ Fixable Viability Kit (BioLegend) (see **Table 4**)
- Distilled water (Gibco)
- FlowMi cell strainers 70um for 1000μL tips (BelART)

**Equipment**

- Hot water bath (37°C)
- Centrifuge
- Nexcelom Auto 2000 Cell Viability Counter

*Thawing Medium*

- 10% Heat-inactivated FBS
- 0.9% Penicillin-Streptomycin
- 1% HEPES
- 0.9% L-Glutamine
- 17.5ng/mL DNase I
- RPMI 1640 with phenol red.
- *Note: The media was warmed in a 37°C water bath before the start of the thawing process*.

*Wash buffers:*

- 2% heat-inactivated FBS in PBS (prepared previous and same day)
- 1X BD Perm/Wash™ Buffer (prepared same day)

*Fixation buffer:*

- 1X Stabilizing Fixative (prepared same day)
- Used at 300μL per test

*Antibody cocktail 1: Viability dye (prepared same day)*

- Zombie Yellow™ Fixable Viability Kit (BioLegend) dye prepared according to manufacturer’s instructions
- 1X PBS to create a working solution at used at 100μL per test with 1:300 Zombie Yellow dye in PBS (0.33μL dye per 100μL volume, see **Table 4**)

*Antibody cocktail 2: iNKT cocktail (prepared same day)*

- Vα24 PE (clone C15)
- áGC-loaded CD1d tetramer BV421
- 1X PBS added to antibody and tetramer for a total (cocktail) volume of 50μL per test

*Antibody cocktail 3: Chemokine receptor cocktail #1 (prepared previous day)*

- CCR5 BUV737 (clone 3A9)
- CXCR3 PE-Cy7 (clone G025H7)
- 1X PBS to antibodies for a total (cocktail) volume of 50μL per test

*Antibody cocktail 4: Chemokine receptor cocktail #2 (prepared previous day)*

- CCR4 BV510 (clone L291H4)
- CCR6 BV650 (clone G034E3)
- 10uL of Brilliant Stain Buffer Plus added to antibodies, followed by PBS added for a total (cocktail) volume of 50μL per test

*Antibody cocktail 5: Surface antibody cocktails (prepared previous day)*

- Antibodies for surface staining (see **Table 4 and Figure 3**)
- 10uL of Brilliant Stain Buffer Plus added to antibodies, followed by PBS added for a total (cocktail) volume of 50μL per test

*Antibody cocktail 6: Intracellular antibody cocktail (prepared previous day)*

- Antibodies for intracellular staining (Granzyme B and Ki67; see **Table 4 and Figure 3**)
- 1X Perm/Wash Buffer added to antibodies for a total (cocktail) volume of 50μL per test

### Protocol

#### Cell preparation

1. Transfer cryovials from liquid nitrogen to a −80°C freezer and store overnight.
2. On the day of the experiment, move cryovials from −80°C to ice chest with with dry ice and thaw in small batches of 3-4 vials.
3. Thaw each cryovial in a 37°C water bath, swirling gently until only a small ice crystal remains. Immediately transfer the closed vial to wet ice.
4. Open the cryovial and add 1mL of pre-warmed thawing medium dropwise directly into it.
5. Transfer the cell suspension slowly into a 15mL conical tube prefilled with 4mL thawing medium.
6. Bring the volume to 15mL with thawing medium.
7. Centrifuge at 350 x *g* for 10 minutes at 4°C.
8. Decant the supernatant and gently resuspend the cell pellet in the residual volume using a 200μL pipette.
9. Add 3 mL of 2% FBS/PBS, mix gently, and centrifuge again at 350 x *g* for 10 minutes at 4°C.
10. Decant the supernatant and resuspend the pellet in 2 mL of 2% FBS/PBS.
11. Visually inspect samples for clumps and filter as needed using FlowMi cell strainers 70μm for 1000μL tips.
12. Determine live leukocyte counts using a Nexcelom Auto 2000 with AO/PI staining (1:1 cell suspension to dye).

#### Cell staining (all steps are performed in the dark and tap vortexed 5x after each reagent is added)

1. Use 1-3 × 10⁶ live leukocytes per sample.
2. Add 3mL of 1X PBS to wash and remove protein in suspension.
3. Centrifuge cells at 350 x g for 5 minutes and decant the supernatant.
4. Resuspend the pellet by gentle pipetting.
5. Add viability dye in PBS (100uL/test at 1:300) and incubate for 20 minutes at room temperature.
6. Wash once with 2% FBS/PBS and centrifuge at 350 x *g* for 5 minutes.
7. Add the iNKT staining cocktail (Vα24-PE and CD1d-tetramer-BV421 in 50μL PBS) and incubate for 20 minutes at room temperature.
8. Add a 50μL per test of anti-CCR5-BUV737, anti-CXCR3-PE-Cy7 in PBS directly to the tube without washing, and incubate for 20 minutes at 37°C.
9. Add a 50μL per test cocktail of anti-CCR4-BV510, anti-CCR6-BV650, Brilliant Stain Buffer Plus in PBS and incubate for 20 minutes at 37°C.
10. Add the remaining surface antibody cocktail at 50μL per test in 2% FBS/PBS and incubate for 20 minutes at room temperature.
11. Wash cells with 3mL of 2% FBS/PBS.
12. Add 200μL BD Cytofix/Cytoperm™ and incubate for 20 minutes at room temperature.
13. Wash with BD Perm/Wash™ buffer (3mL for 5 minutes at 350 x *g*).
14. Add 50μL per test of intracellular antibody cocktail prepared in BD Perm/Wash™ buffer and incubate for 20 minutes at room temperature.
15. Wash once with BD Perm/Wash™ buffer (3mL for 5 minutes at 350 x *g*).
16. Fix cells in 300μL 1x BD Stabilizing Fixative and acquire samples after 1 hour of biohazard inactivation or store at 4°C overnight.
17. Visually inspect samples for clumps, and filter samples using FlowMi cell strainers 70um for 1000μL tips prior to acquisition if clumps are present.

#### Flow cytometry acquisition and analysis

1. Collect at least 10,000 events for single stained controls.
2. Perform initial compensation at the cytometer for real-time quality control of compensation.
3. Acquire samples on a BD FACSymphony A5 (see MIFlowCyt for instrument setup and quality control details).
4. Acquire cell samples at no more than 5,000 events/second using optimized voltages.
5. Re-calculate compensation using traditional compensation in flow cytometry software (Flowjo v10.10).
6. Perform all final gating analyses in flow cytometry software.

### Panel Design and Optimization

#### Overview and Design Objectives

This 28-color flow cytometry panel was developed and optimized on a BD FACSymphony A5 instrument (**Table 3**) to interrogate classical and non-classical T lymphocytes in decidual leukocytes and PBMC from rhesus macaques. The panel resolves phenotypic and functional states of T lymphocytes, including memory differentiation, tissue residency, chemokine receptor expression, activation, exhaustion, cytotoxic potential, and proliferation. Simultaneously, it provides sufficient resolution of B and NK cell subsets to confirm expected frequencies and assess peripheral blood contamination. This panel is complemented by an 18-color panel for deep characterization of NK and myeloid cells in decidua samples (1).

The panel was specifically configured for a finite archive of cryopreserved decidual leukocytes isolated from the maternal-fetal interface and time-matched maternal PBMC. This enables comparisons of protective phenotypes in the peripheral blood and the maternal-fetal immune interface against congenital infections. Since decidual material was limited and irreplaceable, early panel drafting, fluorochrome allocation, and antibody titrations were carried out primarily using PBMC and other available tissues such as lymph nodes, reserving decidual leukocytes for late-stage testing and final validation. Choice of sample type to evaluate a stain was carefully selected considering the biology of each marker. End-stage panel evaluation was carried out on both PBMC and decidual leukocytes. The resulting panel provides a balanced, high-parameter conventional configuration that can be applied to both decidua (**Figure 1**) and PBMC (**Figure 2**) on the same platform without modifying instrument settings or antibody titers.

#### Marker and Clone Selection Strategy

A review of the human literature suggested that immune cells relevant to congenital infections in the decidua include αβ T cells, γδ T cells, iNKT cells, tissue-resident memory (Trm) T cells, T regulatory (Treg) cells, memory T cell subsets, activation and proliferation markers, chemokine receptors, B cells, NK cells, and myeloid antigen-presenting cells (29, 30, 47–50). Since this panel was designed to support evaluation of vaccine-induced T cell responses, we focused on T lymphocyte markers. The final set of markers and reagent information is outlined in **Table 4**.

Antibody selection for non-human primate studies presents unique challenges. Most antibodies used in non-human primate flow cytometry analysis are developed for humans and only incidentally cross-react with rhesus macaques. As a result, clone selection required literature review, consultation of the Nonhuman Primate Reagent Resource (NHPRR), prior experience, and empirical testing of clones without established reactivity in rhesus macaques. In many instances, vendor-reported clonal reactivity to non-human primates lacks supporting data and is unreliable. To validate antibodies lacking prior characterization, candidate clones were screened by parallel staining of rhesus and human PBMC, titrated across relevant tissues, and evaluated by positive-negative discrimination and expected subset frequencies. Only clones demonstrating consistent cross-reactivity and biologically plausible patterns were included.

As an example, the pan-γδ TCR clones 11F2 and B1 both cross-react with humans, but in rhesus macaques, clone 11F2 failed to cross-react entirely and only clone B1 produced expected pan-γδ TCR staining patterns in (**Figure 5**). Similarly, clones B6, REA771, and IMMU389 targeting Vδ2 failed to cross-react in rhesus macaques, whereas clone 15D performed reliably and was ultimately custom conjugated to FITC. In some cases, such as CD57, no cross-reactive reagents were identified.

**Figure 5.**
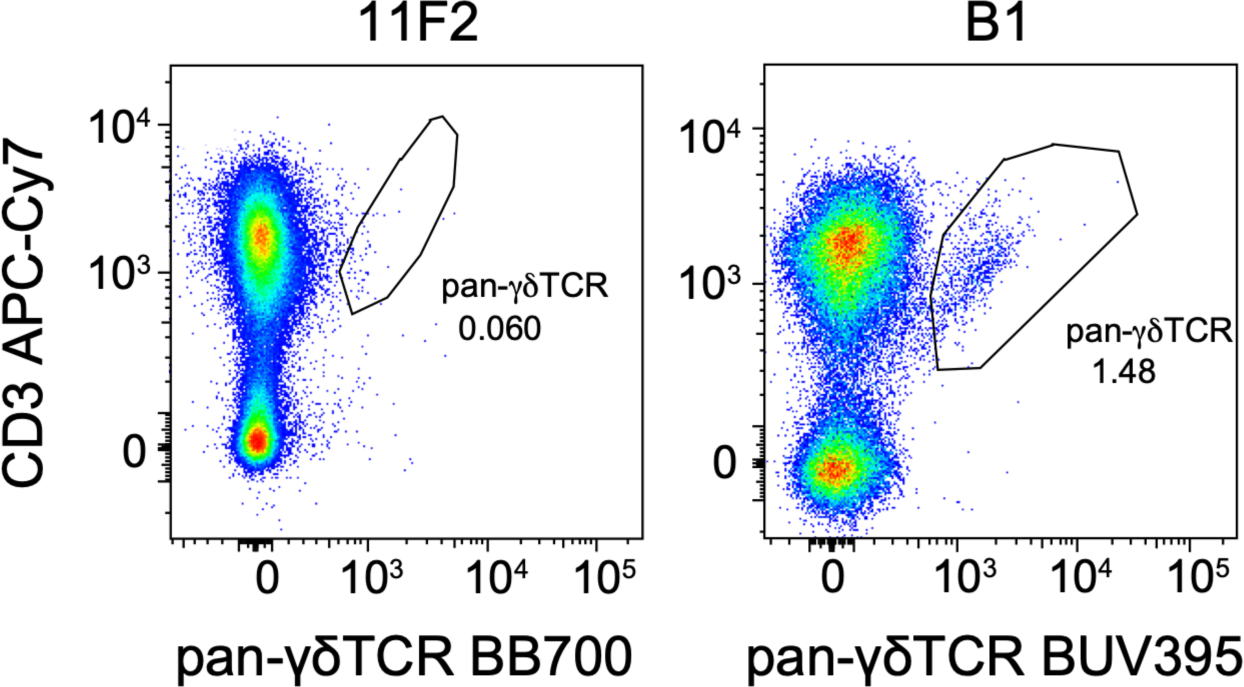
Pan-γδ TCR clone testing for cross-reactivity. After gating on live, singlet leukocytes, γδ T cells were stained with clone 11F2 (left; fluor BB700) or clone B1 (right; fluor BUV395). Expression of pan-γδ TCR is visualized by plotting against CD3 in mesenteric lymph nodes from rhesus macaques.

CX3CR1 presented a more subtle and difficult issue, exemplifying the requirement to validate clones via expected staining patterns rather than simple positive-negative cross-reactivity. The commonly used clone 2A9-1, available in numerous fluor conjugates, demonstrated reliable and clear resolution of positive and negative events across cell types. However, upon deeper analysis, we found that it stained myeloid, B, and CD8+ T cells at similar levels, a pattern inconsistent with published human data (**Figure 6**) (16, 51). Without a strong understanding of the expected biology, these results could easily be misinterpreted as successful cross-reactivity simply because the clone generated clear positive and negative populations. We speculate that clone 2A9-1 may recognize a different antigen altogether or cross-react with an epitope shared among related chemokine receptors. In contrast, clone K0124E1 showed the expected staining pattern but was only commercially available in PE. Therefore, we purchased a custom Alexa Fluor 647 conjugate of clone K0124E1.

**Figure 6.**
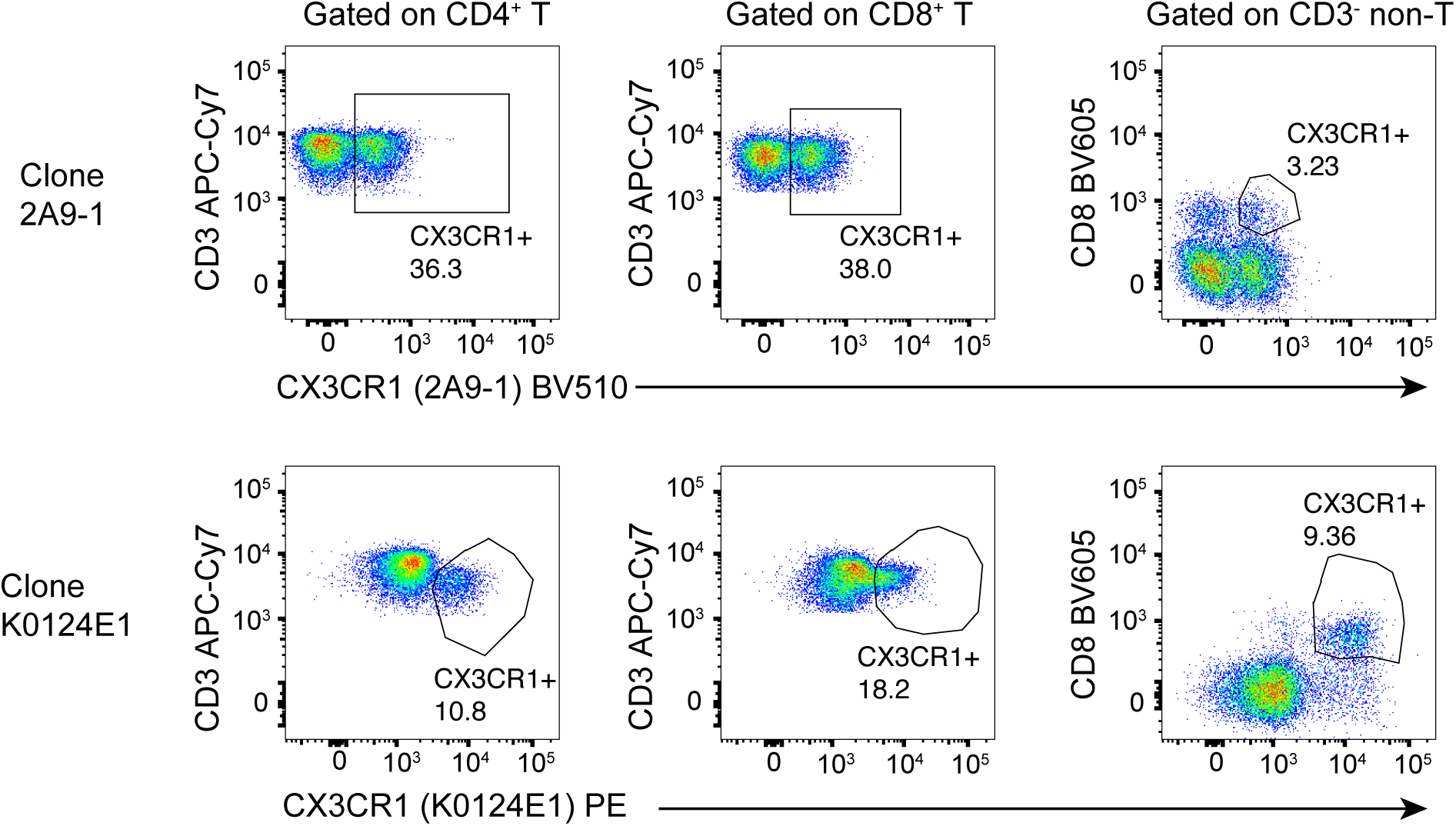
Staining patterns in CX3CR1 vary by clone. Representative flow cytometry dot plots showing the expression of CX3CR1 on CD4SP^+^ T lymphocytes, CD8^+^ T lymphocytes, and CD3^-^ CD8^+^ non-T lymphocytes using clone 2A9-1 (top row) or K0124E1 (bottom row). Data were collected from one healthy animal and using whole blood as the sample type.

Reagents that were evaluated but not included in the final panel are shown in **Figure 7**. The reasons for omitting these reagents were primarily due to issues with cross-reactivity (n=13), good resolution but were reassigned to a different panel or reagent due to panel constraints (n=8), poor resolution (n=8), unexpected staining patterns (n=7), excessive spreading error (n=4) or excessive background (n=1).

**Figure 7.**
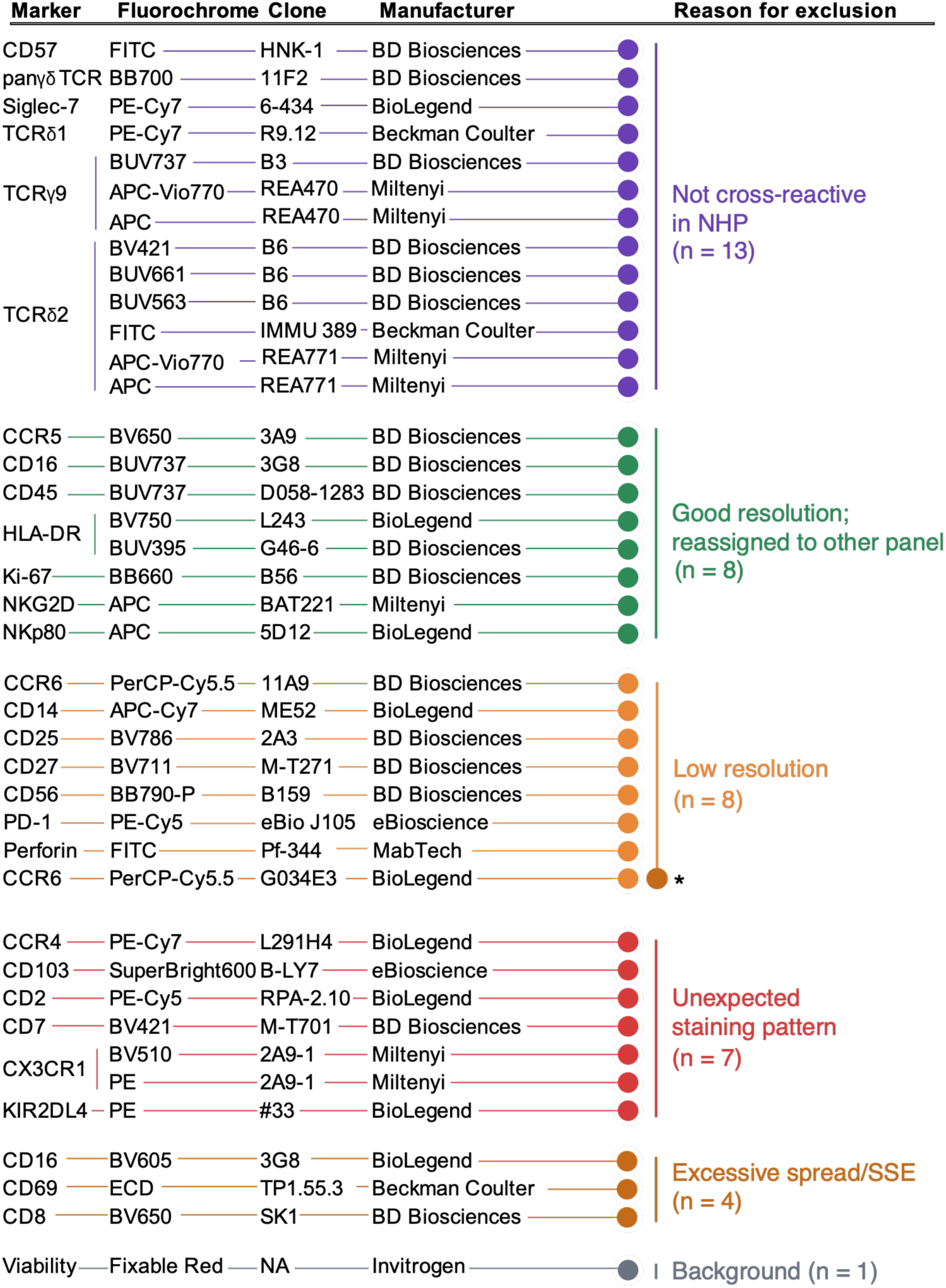
Antibodies evaluated during panel development but excluded from the final panel. Antibodies tested during panel optimization are grouped according to the primary reason for exclusion, including lack of cross-reactivity in NHP, low marker resolution, unexpected staining patterns, excessive spreading error (SSE), or high background staining. Antibodies with acceptable performance that were reassigned to alternative panels or reagents due to panel constraints are also shown. Manufacturer, clone, and fluorochrome information are provided for each antibody. *Indicates a secondary reason for exclusion (excessive spreading error).

#### Fluorochrome and Detector Selection

Fluorochrome assignment was guided by antigen density, clone availability, commercially available conjugations, and the need to minimize spillover and spreading error in a 28-color configuration on the BD FACSymphony A5. High-density antigens and lineage markers were placed on dimmer or moderate fluorochromes, whereas low-density or functionally critical markers (*e.g.,* chemokine receptors, Treg markers, and γδ T cell markers) were assigned to brighter fluorochromes. For example, CD25 was more clearly resolved on BUV661 than BV786 (**Figure 8**). Chemokine receptors required particular attention due to low antigen density, co-expression patterns, and limited cross-reactive fluorophore options.

**Figure 8.**
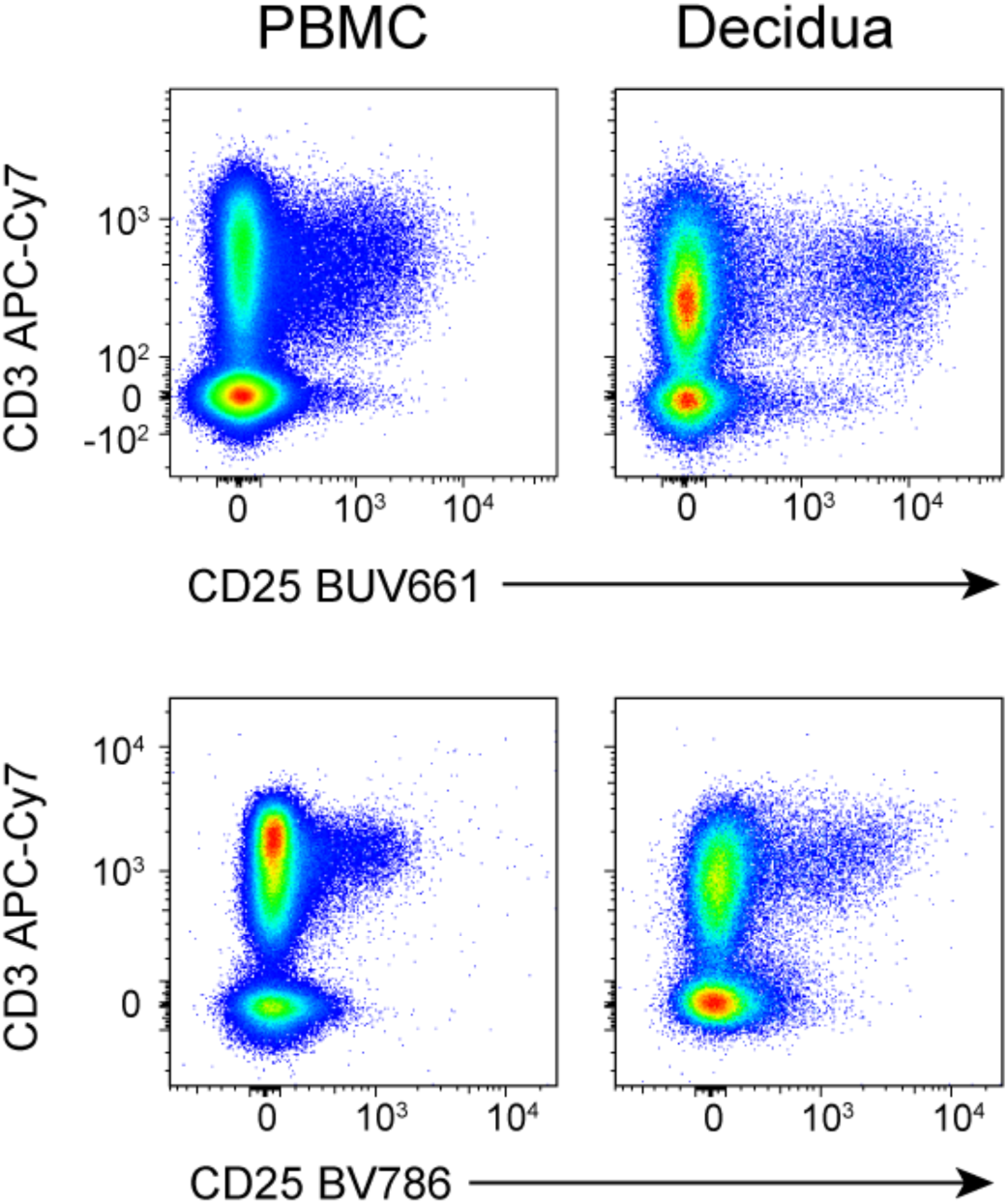
Fluorochrome optimization for CD25. PBMC and decidual leukocytes are fully stained with either CD25 BUV661 (top) or BV786 (bottom) with the same clone (2A3).

At the time of this panel design, few commercially available rhesus-cross-reactive clones were conjugated to the newer fluorophores needed to occupy novel spectral spaces on a Symphony A5. For example, only three rhesus-cross-reactive clones relevant to this panel (CD20, CD56, HLA-DR) were available conjugated to BV570. For pan-γδ TCR, an alternative cross-reactive clone 5A6.E9 may further improve detection of a subset of γδT cells, but was only available in the fluorophores PE, FITC, and TRI-COLOR (also called PE-Cy5), which were committed for other key markers (Vα24 and Vδ2). This severely restricted panel design options in the long wavelength blue, violet, and UV detector-arrays.

Spillover and spreading error (SSE) were evaluated throughout panel development. High-spread interactions were first identified using the Spillover Spreading Matrix (SSM; **Table 5**). Thirteen fluorochrome pairs had SSE > 3, indicating potential resolution issues. Co-expressed markers were avoided on high-spread fluor pairs. In one case, a very high-spread pair (BB630 → BV605; SSM = 13.74) involved markers on distinct lineages (CD28 and CD20), preventing misclassification despite the high SSE. Bright fluorochromes with greater potential for spreading were reserved for low-density markers, whereas dimmer fluorochromes were allocated to high-density markers less sensitive to spread.

**Table 5.** The spillover spread matrix (SSM) calculated in FlowJo using BioLegend compensation beads, with the universal negative control applied for all fluorochromes and amine-reactive positive and negative beads used for the Zombie Yellow viability dye. Thirteen fluorochrome pairs yielded spillover spreading error (SSE) values >3 and are flagged in red. Source conjugates are listed vertically, and detected conjugates are listed horizontally.

|  | AF647 | AF700 | APC-Cy7 | FITC | BB630 | BB660 | BB700 | BB790-P | BUV396 | BUV496 | BUV563 | BUV615 | BUV661 | BUV737 | BUV805 | BV421 | BV510 | BV570 | BV605 | BV650 | BV711 | BV750 | Zombie Yell | PE | PE-CF594 | PE-Cy5-5 | PE-Cy5 | PE-Cy7 |
| --- | --- | --- | --- | --- | --- | --- | --- | --- | --- | --- | --- | --- | --- | --- | --- | --- | --- | --- | --- | --- | --- | --- | --- | --- | --- | --- | --- | --- |
| AF647 | 0.00 | 1.22 | 0.27 | 0.00 | 0.08 | 0.80 | 0.64 | 0.27 | 0.04 | 0.05 | 0.09 | 0.12 | 0.33 | 0.18 | 0.11 | 0.00 | 0.00 | 0.19 | 0.17 | 0.28 | 0.26 | 0.14 | 0.14 | 0.06 | 0.09 | 0.61 | 1.09 | 0.38 |
| AF700 | 0.29 | 0.00 | 0.39 | 0.00 | 0.22 | 0.31 | 1.07 | 0.44 | 0.10 | 0.16 | 0.20 | 0.23 | 0.22 | 0.71 | 0.44 | 0.00 | 0.00 | 0.37 | 0.33 | 0.21 | 0.61 | 0.62 | 0.44 | 0.06 | 0.11 | 0.91 | 0.30 | 0.51 |
| APC-Cy7 | 0.44 | 0.32 | 0.00 | 0.00 | 0.26 | 0.32 | 0.47 | 1.13 | 0.10 | 0.01 | 0.25 | 0.27 | 0.34 | 0.52 | 1.36 | 0.00 | 0.00 | 0.43 | 0.30 | 0.15 | 0.27 | 0.34 | 0.57 | 0.15 | 0.21 | 0.37 | 0.31 | 1.90 |
| FITC | 0.07 | 0.00 | 0.00 | 0.00 | 0.29 | 0.18 | 0.26 | 0.08 | 0.00 | 0.19 | 0.22 | 0.23 | 0.14 | 0.10 | 0.00 | 0.00 | 0.00 | 0.35 | 0.39 | 0.16 | 0.15 | 0.07 | 0.07 | 0.05 | 0.12 | 0.13 | 0.00 | 0.00 |
| BB630 | 0.66 | 0.34 | 0.09 | 2.05 | 0.00 | 1.20 | 1.73 | 0.55 | 0.26 | 0.48 | 0.45 | 3.59 | 0.83 | 0.35 | 0.26 | 0.36 | 0.71 | 2.71 | 13.75 | 1.65 | 0.78 | 0.54 | 0.34 | 0.63 | 4.15 | 0.70 | 1.38 | 0.25 |
| BB660 | 3.16 | 1.02 | 0.22 | 0.00 | 0.40 | 0.00 | 1.96 | 0.62 | 0.05 | 0.07 | 0.16 | 0.24 | 2.03 | 0.62 | 0.35 | 0.00 | 0.00 | 0.57 | 0.59 | 1.64 | 1.13 | 0.86 | 0.50 | 0.06 | 0.19 | 0.51 | 0.92 | 0.23 |
| BB700 | 0.91 | 1.19 | 0.22 | 0.00 | 0.34 | 1.03 | 0.00 | 0.83 | 0.06 | 0.08 | 0.13 | 0.18 | 0.96 | 0.74 | 0.43 | 0.00 | 0.00 | 0.45 | 0.45 | 0.71 | 1.64 | 1.09 | 0.63 | 0.04 | 0.12 | 0.51 | 0.35 | 0.32 |
| BB790-P | 0.16 | 0.14 | 0.22 | 0.00 | 0.23 | 0.26 | 0.44 | 0.00 | 0.13 | 0.11 | 0.18 | 0.22 | 0.19 | 0.35 | 1.29 | 0.00 | 0.00 | 0.37 | 0.37 | 0.18 | 0.24 | 0.32 | 0.80 | 0.07 | 0.15 | 0.13 | 0.06 | 0.29 |
| BUV396 | 0.07 | 0.07 | 0.04 | 0.00 | 0.13 | 0.13 | 0.18 | 0.08 | 0.00 | 0.37 | 0.24 | 0.24 | 0.17 | 0.07 | 3926 | 0.00 | 0.00 | 0.12 | 0.22 | 0.11 | 0.12 | 0.07 | 0.00 | 0.00 | 0.00 | 0.19 | 0.09 | 0.00 |
| BUV496 | 0.13 | 0.10 | 0.05 | 0.25 | 0.32 | 0.13 | 0.24 | 0.00 | 0.29 | 0.00 | 0.82 | 0.76 | 0.51 | 0.26 | 0.17 | 0.00 | 0.77 | 0.70 | 0.49 | 0.18 | 0.15 | 0.14 | 0.07 | 0.14 | 0.31 | 0.09 | 0.09 | 0.00 |
| BUV563 | 0.39 | 0.05 | 0.03 | 0.33 | 0.90 | 0.39 | 0.42 | 0.19 | 0.42 | 0.24 | 0.00 | 1.59 | 0.94 | 0.34 | 0.22 | 0.00 | 0.00 | 1.48 | 1.19 | 0.28 | 0.21 | 0.14 | 0.10 | 1.00 | 0.95 | 0.26 | 0.45 | 0.12 |
| BUV615 | 0.63 | 0.20 | 0.12 | 0.26 | 3.35 | 0.55 | 0.68 | 0.24 | 1.43 | 0.35 | 0.78 | 0.00 | 1.51 | 0.63 | 0.39 | 0.00 | 0.64 | 1.15 | 2.56 | 0.38 | 0.32 | 0.25 | 0.16 | 0.40 | 3.94 | 0.58 | 0.78 | 0.29 |
| BUV661 | 2.06 | 0.73 | 0.17 | 0.00 | 0.17 | 0.65 | 0.59 | 0.22 | 0.16 | 0.08 | 0.13 | 0.31 | 0.00 | 0.94 | 0.55 | 0.00 | 0.00 | 0.18 | 0.22 | 0.64 | 0.39 | 0.31 | 0.21 | 0.07 | 0.20 | 0.51 | 0.95 | 0.30 |
| BUV737 | 0.13 | 0.64 | 0.26 | 0.00 | 0.10 | 0.33 | 1.09 | 0.59 | 0.25 | 0.11 | 0.11 | 0.14 | 0.23 | 0.00 | 1.36 | 0.00 | 0.00 | 0.11 | 0.16 | 0.10 | 0.39 | 0.66 | 0.45 | 0.00 | 0.05 | 0.25 | 0.10 | 0.23 |
| BUV805 | 0.00 | 0.15 | 0.27 | 0.00 | 0.14 | 0.09 | 0.19 | 0.35 | 0.54 | 0.15 | 0.23 | 0.31 | 0.24 | 0.45 | 0.00 | 0.00 | 0.00 | 0.26 | 0.26 | 0.14 | 0.09 | 0.10 | 0.25 | 0.05 | 0.16 | 0.14 | 0.00 | 0.28 |
| BV421 | 0.00 | 0.00 | 0.00 | 0.00 | 0.10 | 0.07 | 0.12 | 0.13 | 0.00 | 0.34 | 0.18 | 0.22 | 0.13 | 0.08 | 0.08 | 0.00 | 0.00 | 0.39 | 0.27 | 0.11 | 0.12 | 0.11 | 0.08 | 0.00 | 0.07 | 0.11 | 0.00 | 0.00 |
| BV510 | 0.18 | 0.20 | 4289 | 0.00 | 0.45 | 0.13 | 0.22 | 0.00 | 0.13 | 1.09 | 0.79 | 0.92 | 0.64 | 0.33 | 0.24 | 0.00 | 0.00 | 1.86 | 1.34 | 0.53 | 0.48 | 0.40 | 0.26 | 0.18 | 0.42 | 0.10 | 0.12 | 0.00 |
| Zombie Yell | 0.56 | 0.44 | 0.06 | 0.85 | 1.48 | 0.46 | 0.63 | 0.35 | 0.93 | 1.35 | 2.37 | 1.81 | 1.17 | 0.59 | 0.37 | 0.94 | 0.50 | 0.00 | 2.79 | 0.94 | 1.04 | 0.92 | 0.58 | 0.43 | 0.91 | 0.28 | 0.25 | 0.00 |
| BV605 | 0.46 | 0.37 | 0.07 | 0.00 | 1.31 | 0.56 | 0.70 | 0.28 | 0.14 | 0.25 | 0.78 | 1.71 | 1.21 | 0.53 | 0.39 | 0.00 | 0.29 | 2.15 | 0.00 | 0.89 | 0.92 | 0.74 | 0.49 | 0.45 | 1.29 | 0.56 | 0.69 | 0.26 |
| BV650 | 1.08 | 0.81 | 0.20 | 0.00 | 0.40 | 0.56 | 0.77 | 0.30 | 0.08 | 0.09 | 0.25 | 0.58 | 2.58 | 0.89 | 0.55 | 0.00 | 0.00 | 0.65 | 0.95 | 0.00 | 1.50 | 1.23 | 0.73 | 0.13 | 0.41 | 0.61 | 0.74 | 0.28 |
| BV711 | 0.40 | 1.92 | 0.36 | 0.00 | 0.08 | 0.39 | 1.27 | 0.48 | 0.14 | 0.10 | 0.15 | 0.18 | 0.60 | 2.37 | 1.11 | 0.24 | 0.00 | 0.25 | 0.26 | 0.42 | 0.00 | 1.79 | 1.18 | 0.05 | 0.10 | 0.58 | 0.22 | 0.35 |
| BV750 | 0.00 | 0.41 | 0.24 | 0.00 | 0.20 | 0.19 | 0.61 | 0.44 | 0.16 | 0.10 | 0.25 | 0.27 | 0.17 | 2.47 | 1.44 | 0.00 | 0.00 | 0.37 | 0.33 | 0.20 | 0.44 | 0.00 | 1.39 | 0.11 | 0.15 | 0.29 | 0.10 | 0.28 |
| BV786 | 0.08 | 0.20 | 0.28 | 0.00 | 0.00 | 0.00 | 0.22 | 0.48 | 0.23 | 0.12 | 0.16 | 0.15 | 0.14 | 0.97 | 2.36 | 1.78 | 0.00 | 0.25 | 0.19 | 0.16 | 0.26 | 0.76 | 0.00 | 0.05 | 0.00 | 0.17 | 0.00 | 0.31 |
| PE | 0.52 | 0.18 | 0.00 | 0.00 | 1.82 | 1.13 | 1.10 | 0.42 | 0.29 | 0.46 | 2.66 | 1.19 | 0.69 | 0.37 | 0.19 | 0.00 | 0.00 | 4.03 | 2.67 | 0.67 | 0.59 | 0.39 | 0.22 | 0.00 | 1.51 | 0.67 | 0.88 | 0.34 |
| PE-CF594 | 0.67 | 0.26 | 0.08 | 0.00 | 3.96 | 1.45 | 1.65 | 0.58 | 0.16 | 0.22 | 0.46 | 2.37 | 0.75 | 0.31 | 0.22 | 0.00 | 0.59 | 1.89 | 3.38 | 0.78 | 0.66 | 0.43 | 0.25 | 0.50 | 0.00 | 1.00 | 1.57 | 0.50 |
| PE-Cy5-5 | 0.52 | 0.69 | 0.16 | 0.00 | 0.30 | 1.15 | 3.61 | 1.01 | 0.06 | 0.08 | 0.25 | 0.17 | 0.67 | 0.64 | 0.36 | 0.00 | 0.00 | 0.54 | 0.40 | 0.46 | 1.31 | 0.73 | 0.44 | 0.31 | 0.31 | 0.00 | 0.94 | 0.94 |
| PE-Cy5 | 1.65 | 0.59 | 0.15 | 0.00 | 0.17 | 2.29 | 2.15 | 0.70 | 0.04 | 0.04 | 0.10 | 0.11 | 1.64 | 0.35 | 0.23 | 0.00 | 0.00 | 0.31 | 0.35 | 1.18 | 0.79 | 0.50 | 0.26 | 0.12 | 0.14 | 1.35 | 0.00 | 0.65 |
| PE-Cy7 | 0.05 | 0.06 | 0.23 | 0.00 | 0.15 | 0.13 | 0.27 | 3.11 | 0.09 | 0.03 | 0.11 | 0.10 | 0.06 | 0.28 | 0.81 | 0.00 | 0.00 | 0.21 | 0.17 | 0.05 | 0.08 | 0.13 | 0.66 | 0.10 | 0.13 | 0.18 | 0.10 | 0.00 |

In addition to SSM-based evaluation, an important stage included NxN plots gated on live leukocytes that were used throughout panel development to manually assess biologically relevant staining patterns in decidua and PBMC, verify compensation accuracy, and identify unacceptable spreading (**Figure 9**).

**Figure 9.**
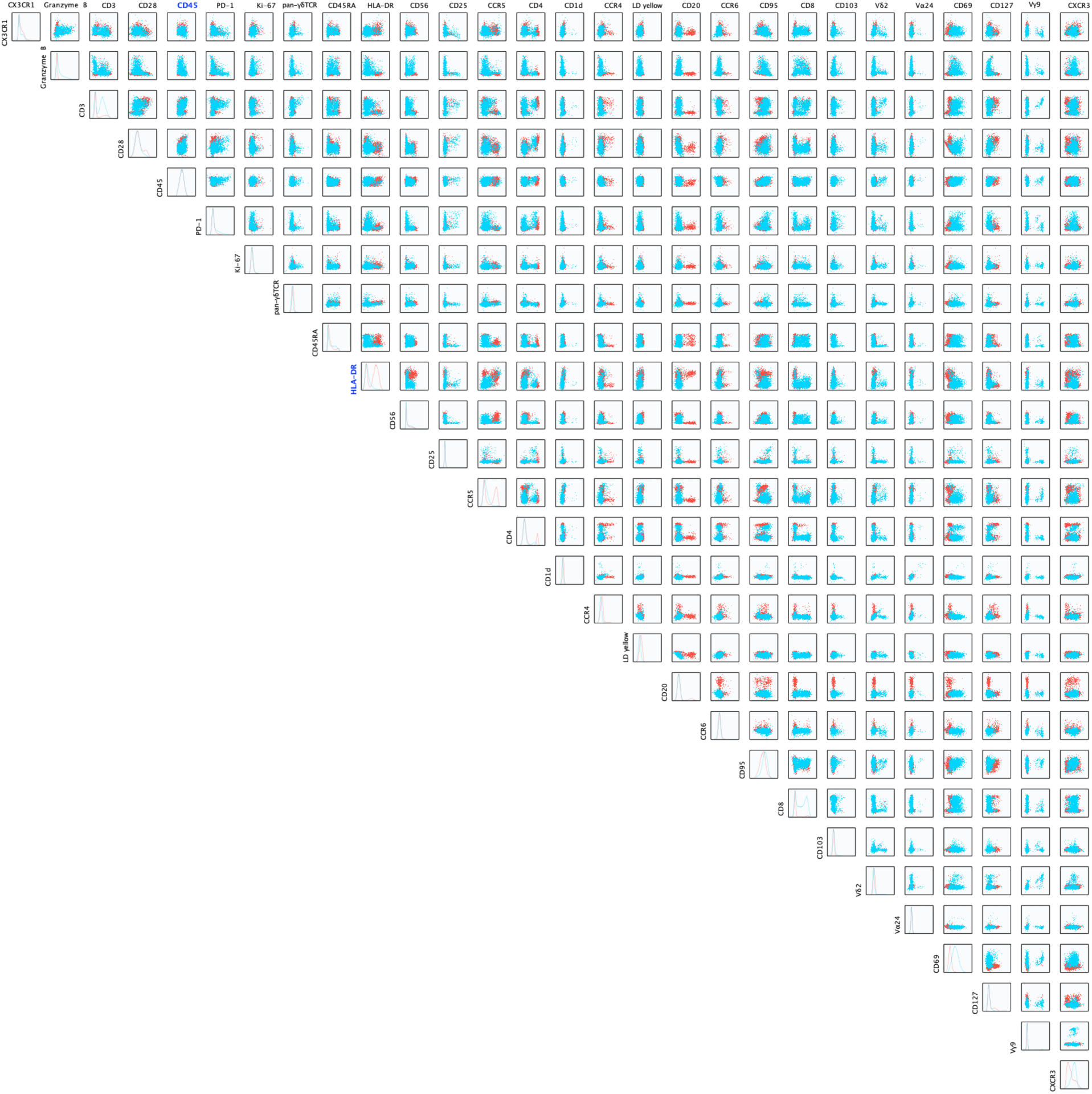
NxN plot gated on time/single cells/CD45+/live/lymph of decidua (blue) and PBMC (red). NxN plots enable overview assessment of spread and biological staining patterns for both sample types at once.

Importantly, we identified a dim false positive population in the PE channel that resulted from incomplete compensation of the BUV563 parameter. This occurred while using Spherotech COMPtrol compensation beads, which were dimmer with BUV563 than the corresponding samples. Since BUV563 spills heavily into PE, the dimness of the compensation control with BUV563 caused incomplete spillover correction and created a false positive effect in PE. Using a brighter bead control (BioLegend compensation beads) reduced the incidence of PE-positive events into populations such as Vα24^+^ iNKTs (**Figure 10**). This issue was more pronounced in decidua which has more CD3^-^ HLA-DR^+^ bright events than PBMC. These results were consistent with the requirement that compensation controls be as bright or brighter than the stained samples, and the BioLegend compensation controls were used in the final panel analysis. In addition, the amount of BUV563 HLA-DR was reduced at a sub-saturating titer to reduce SSE into PE. The reduction in titer still resulted in sufficient resolution of HLA-DR positive events for the objectives of this panel.

**Figure 10.**
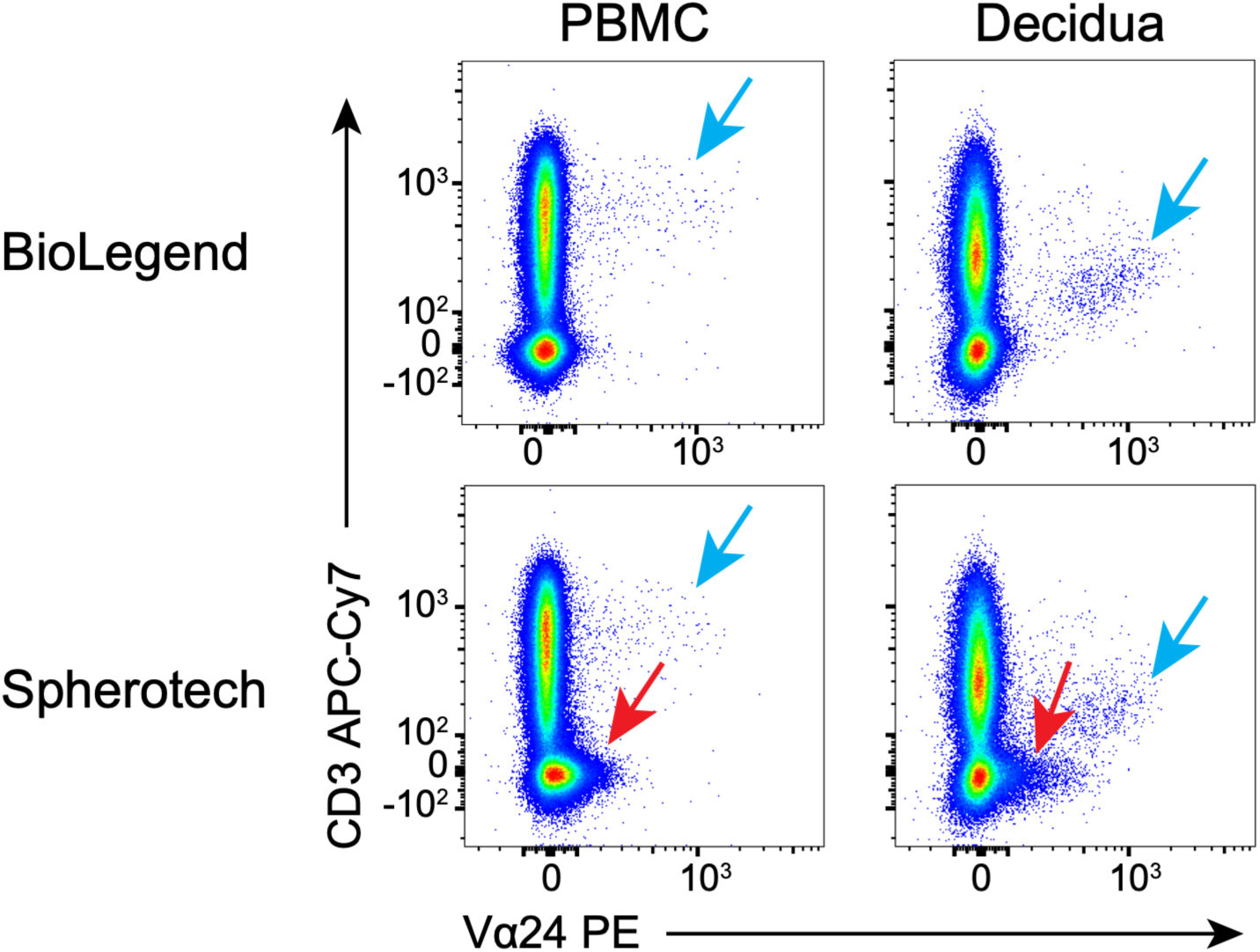
Spreading of PE Vα24 using different compensation beads. Dot plot representations of PBMC and decidual leukocytes stained and gated on live/single/CD45^+^ lymphocytes followed by visualization of CD3 APC-Cy7 vs Vα24 PE. Plots show data from the same sample after compensating with BioLegend beads (top row) or Spherotech (bottom row) COMPtrol beads. Red arrows highlight events that are undercompensated from other channels when using Spherotech beads, which is not observed when using BioLegend beads. Blue arrows highlight the true positive Vα24+ CD3+ T cells.

Early panel drafts were used to identify problematic fluor combinations before the full reagent set was assembled. Two initial “skeleton” panels, one based on idealized spillover modeling, and one based on known rhesus-compatible pairings, were tested to identify fluorochromes that performed reliably across tissues and to guide refinements. Subsequent modifications included relocating bright fluorochromes (*e.g.,* BUV737, BUV661, BUV615) to markers requiring greater sensitivity and reassigning markers that experienced excess spread (*e.g.,* CD8 on BV650, CD25 on BV786) to more appropriate channels.

Final fluor-marker selections reflect 1) empirical testing of clone-fluor combinations, 2) minimization of conflicts among co-expressed markers, and 3) balanced use of available detectors. A summary of panel iterations is provided in **Table 6**.

**Table 6.** Experiment iterations of panel development.

| Panel A iteration | 1 (8-color) | 2 (12-color) | 3 (21-color) | 4 (21-color) | 5 (24-color) | 6 (25-color) | 7 (25-color) | 8 (28-color FINAL) |
| --- | --- | --- | --- | --- | --- | --- | --- | --- |
| Sample type | Spleen | Spleen | MLN and Mesfat | MLN but different L/D | Fresh PBMC and decidua | PBMC different L/D and CCR4 | Fresh PBMC and decidua | PBMC and decidua |
| FITC | Perforin | Perforin | Perforin | Perforin | Perforin | Perforin |  | Vδ2 |
| BB630 |  |  |  |  | CD28 | CD28 | CD28 | CD28 |
| BB660 |  |  |  |  | CD45 | CD45 | CD45 | CD45 |
| BB700 |  |  | pan-γδTCR | pan-γδTCR |  |  | PD-1 | PD-1 |
| BB790 |  |  |  |  | Ki-67 | Ki-67 | Ki-67 | Ki-67 |
| BV421 | CD1d | CD1d | CD1d | CD1d | CD1d | CD1d | CD1d | CD1d |
| BV510 | L/D aqua | L/D aqua | L/D aqua |  | L/D aqua | CCR4 | CCR4 | CCR4 |
| BV570 |  |  |  | L/D yellow |  | L/D yellow | LD yellow | LD yellow |
| BV605 |  | CD16 | CD16 | CD16 | CD16 | CD16 | CD20 | CD20 |
| BV650 |  | CCR6 | CCR6 | CCR6 | CCR6 | CCR6 | CCR6 | CCR6 |
| BV711 | CD95 | CD95 | CD95 | CD95 | CD95 | CD95 | CD95 | CD95 |
| BV750 |  | CD8 | CD8 | CD8 | CD8 | CD8 | CD8 | CD8 |
| SB780 |  |  |  |  |  |  | CD103 | CD103 |
| BUV396 |  |  |  |  | HLA-DR | HLA-DR | pan-γδTCR | pan-γδTCR |
| BUV496 |  |  | CD45RA | CD45RA | CD45RA | CD45RA | CD45RA | CD45RA |
| BUV563 |  |  | Vδ2 | Vδ2 |  |  |  | HLA-DR |
| BUV615 |  |  | CD56 | CD56 | CD56 | CD56 | CD56 | CD56 |
| BUV661 |  |  | CD25 | CD25 | CD25 | CD25 | CD25 | CD25 |
| BUV805 |  |  | CD4 | CD4 | CD4 | CD4 | CD4 | CD4 |
| BUV737 |  |  | CCR5 | CCR5 | CCR5 | CCR5 | CCR5 | CCR5 |
| PE | Vα24 | Vα24 | Vα24 | Vα24 | Vα24 | Vα24 | Vα24 | Vα24 |
| PE-CF594 | CD69 | CD69 | CD69 | CD69 | CD69 | CD69 | CD69 | CD69 |
| PE-Cy5 |  |  | PD-1 | PD-1 | PD-1 | PD-1 | Vγ9 | Vγ9 |
| PE-Cy5.5 |  |  | CD127 | CD127 | CD127 | CD127 | CD127 | CD127 |
| PE-Cy7 |  | CXCR3 | CXCR3 | CXCR3 | CXCR3 | CXCR3 | CXCR3 | CXCR3 |
| AF647 |  |  |  |  | NKG2D (APC) | NKG2D (APC) |  | CX3CR1 |
| AF700 | Granzyme B | Granzyme B | Granzyme B | Granzyme B | Granzyme B | Granzyme B | Granzyme B | Granzyme B |
| APC-Cy7 | CD3 | CD3 | CD3 | CD3 | CD3 | CD3 | CD3 | CD3 |

#### Antibody staining and validation

All antibodies were titrated using PBMC or tissue depending on antigen abundance (**Table 4**; **Figure 11**). Viability dye was included with all stains, and compensation was performed for each titration analysis. Two-fold serial dilutions for a total of five dilutions were performed for each antibody, with a starting dilution volume of 2X the manufacturer recommendation and a final dilution volume of 0.125X the manufacturer recommendation. Antibodies were prepared in 50μL cocktail test volume and added after cells were stained for live/dead, washed, and decanted. The viability dye was titrated in both PBMC and spleen to identify a concentration suitable across sample types. Tissue-specific differences were subtle and an acceptable concentration (1:300) was selected to work for both tissues. Primary markers were regularly co-stained with a limited number (1–4) of additional markers to maximize resolution or validate reagents. For example, the inclusion of live/dead, CD3, and Vα24 antibody with the CD1d-tetramer titration and CD1d-tetramer negative control assured specificity. Titers were selected based on visual clarity of positive/negative discrimination and limited background of the negative populations. After selection of initial titers, partial and full test panels were used to confirm expected subset frequencies and high resolution when mixed with other antibodies and after implementation of the sequential staining workflow. The final selected titers are indicated in **Figure 11** and **Table 4**.

**Figure 11.**
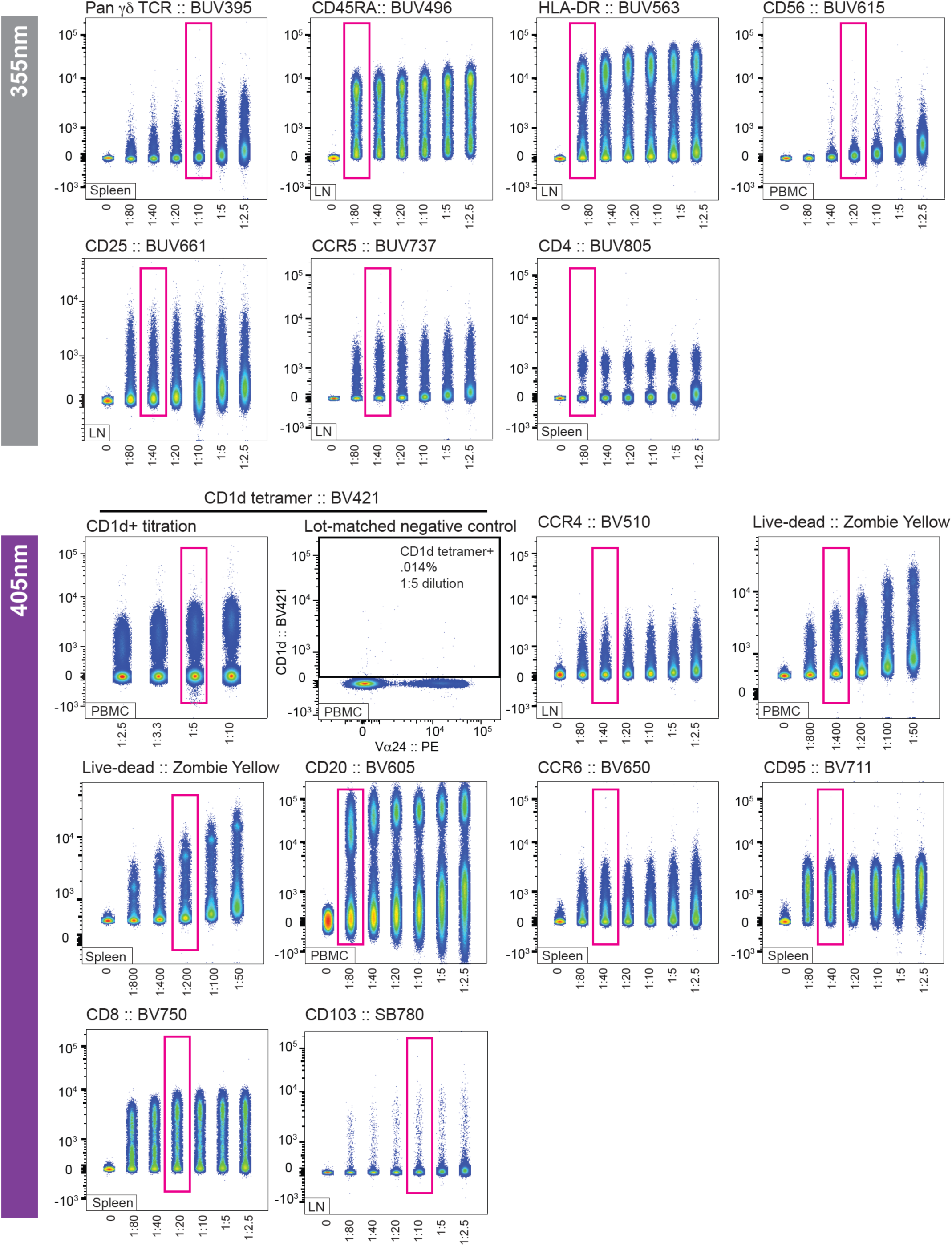
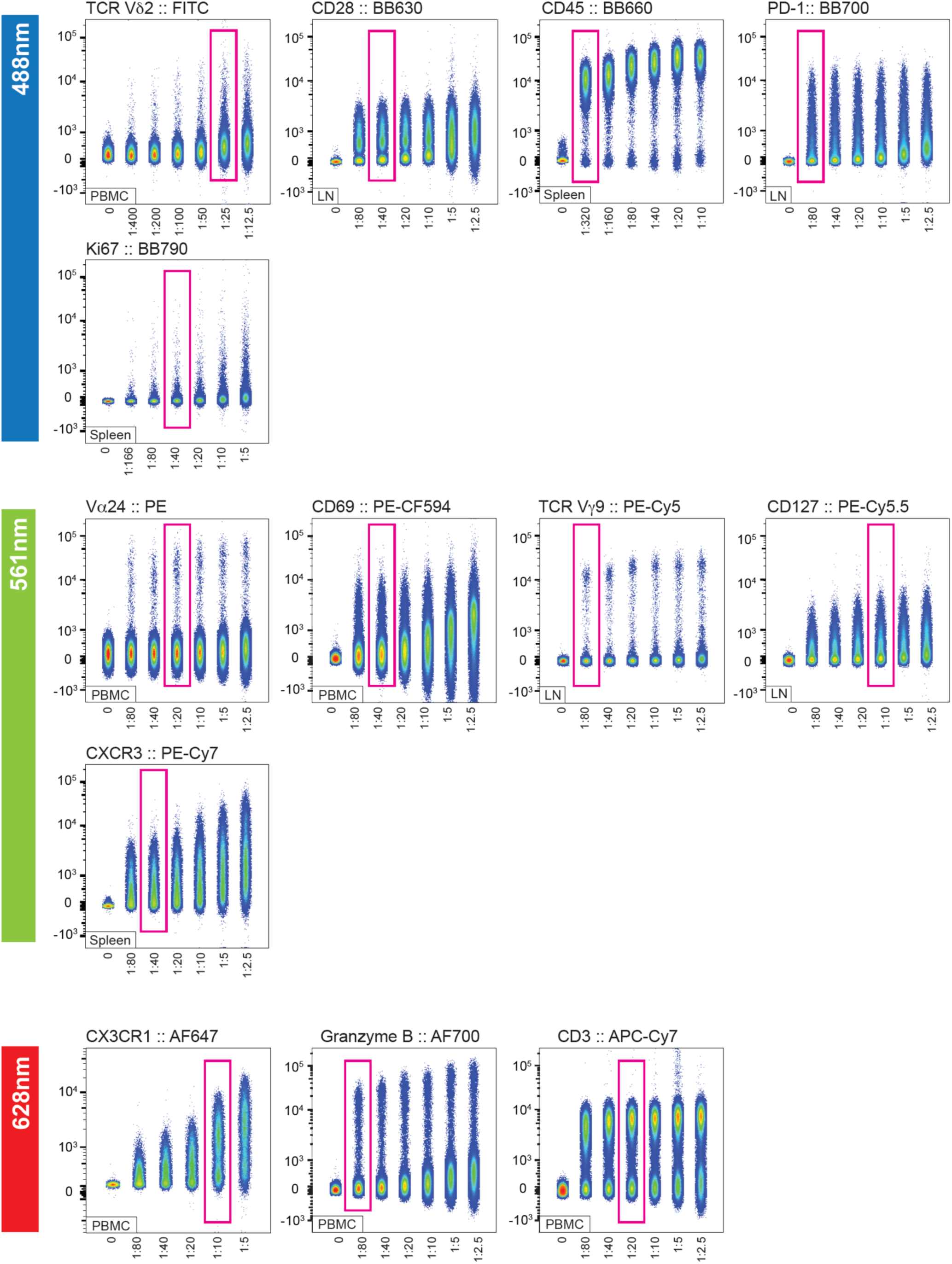
Antibody titrations. Individual titration FCS files were gated on single/live cells (except viability dye titration, which was gated only on singlets) and then concatenated into one master FCS file for each antibody. Master concatenated FCS files are shown here and were analyzed by displaying the Sample ID on the x-axis (*i.e.,* one sample/FCS file per ID), and the titrated antibody on the y-axis. Pink boxes indicate the final selected antibody dilution. The tissue on which the antibody was titrated is indicated in the bottom left of each plot. An additional negative control that includes several co-stains (live/dead, CD3, and Vα24) is added alongside the CD1d titration to assess the background of BV421 CD1d.

Sequential staining was required because several key markers exhibit known reduced resolution when stained together. Tetramer binding can be reduced if stained in combination with reagents that may interfere (*i.e.,* CD4 and CD8 antibodies) while chemokine and cytokine receptors both have mobile transmembrane domains and also undergo regular recycling (52–54). Detection of both is improved when stained prior to the remaining surface antibodies, and chemokine/cytokine receptor staining is often improved at elevated temperatures (52). A priori laboratory testing with staining nonhuman primate mucosal tissues guided the decision to stain CCR5/CXCR3 and CCR4/CCR6 in two independent antibody cocktails. The amine-reactive dye to detect dead cells at optimal resolution requires that cells are stained and washed with 1XPBS to remove amine-rich proteins in suspension, whereas the remaining stains are performed in cells resuspended in 2% fetal bovine serum (FBS) in PBS. The resulting staining sequence was finalized as follows: samples were first stained with the viability dye, followed by Vα24 and CD1d tetramer, then CCR5 and CXCR3, followed by CCR4 and CCR6, the remaining surface marker cocktail, and finally the intracellular markers. Chemokine receptor staining was performed at 37°C, whereas all other staining steps were conducted at room temperature. Following surface staining, cells were fixed and permeabilized prior to intracellular staining. Collectively, this sequential workflow enabled robust detection of chemokine receptors, lineage markers, and intracellular targets within a single high-parameter panel. The sequential staining workflow and antibody cocktail reagents and volumes are illustrated in **Figure 3**, detailed in the protocol, and reported in **Table 4**.

Full-panel staining of PBMC and decidua was used to confirm expected tissue biology. As expected, decidua showed enrichment of NK cells, regulatory lymphocytes, lower frequencies of B cells (**Figures 1-2**), and a CD69^+^CD103^+^ tissue-resident T cells enriched in decidua compared with PBMC which also varied significantly by animal (**Figure 12**). The CD25 expression on CD4SP^+^ T lymphocytes was also brighter in decidua, as expected. In addition, NK cell phenotypes differed between tissues, with CD56^+^ NK cells predominating in decidua and PBMC consisting largely of CD56^-^ NK cells, which is an established pattern in rhesus macaques (1, 55) (**Figures 1-2**).

**Figure 12.**
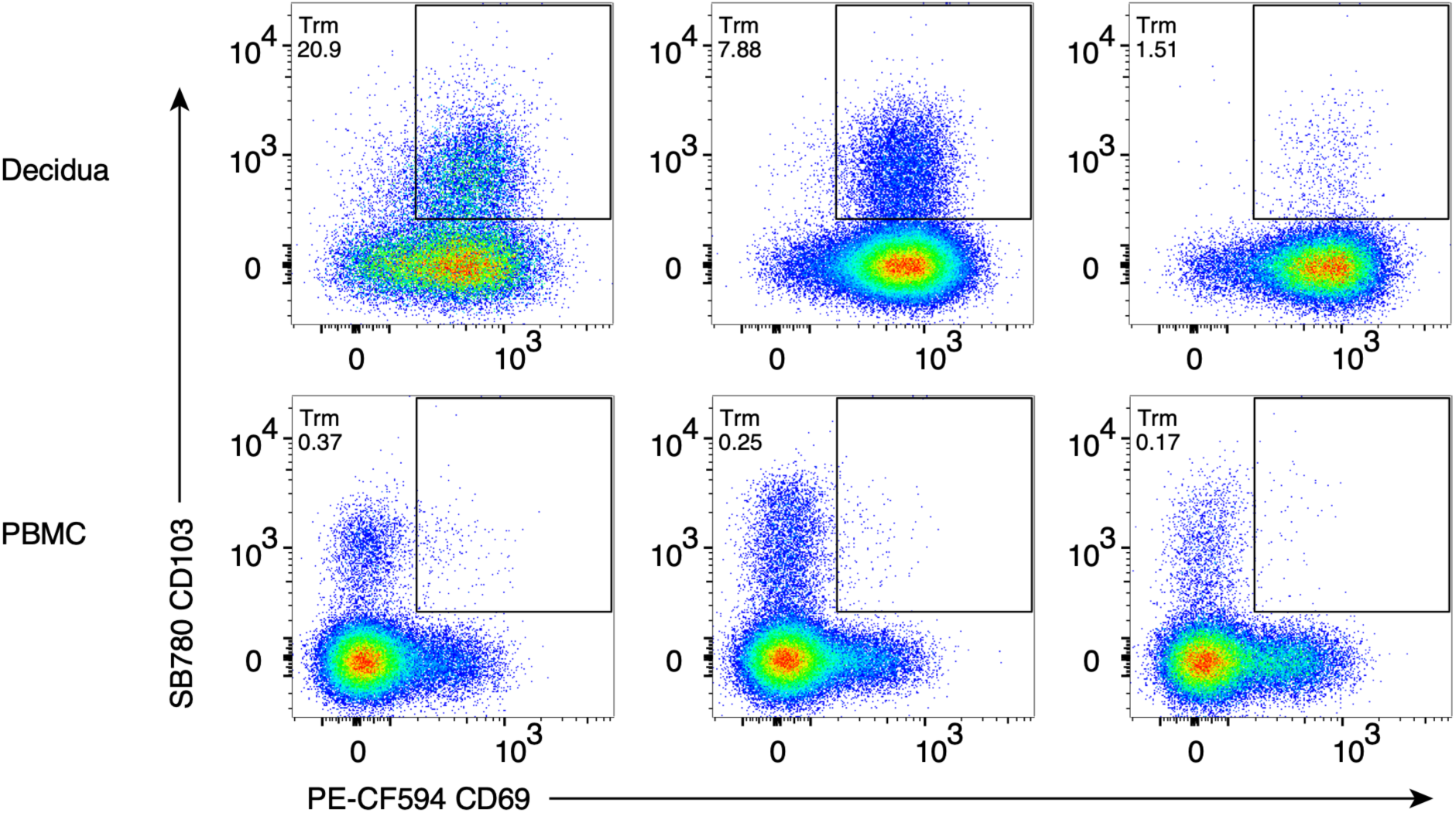
Representative dot plots of Tissue resident memory T cells in PBMC and decidua. CD103 and CD69 expression from three different rhesus macaques (one per column) on memory CD8+ T lymphocytes are displayed. The population has varying expression in decidua (top row) and is not commonly found in circulating cells (bottom row). Cell type frequencies are shown as a percent of total memory CD8^+^ T lymphocytes and are shown in order of high, medium, and low frequency.

Single color compensation controls and a universal negative were made using BioLegend compensation beads for all fluorescent parameters except viability, for which positive and negative Amine Reactive Compensation (ArC) beads were used together. Fluorescence-minus-one (FMO) controls were used to determine gate placement for continuous markers that can be difficult to distinguish from background spread, including PD-1, HLA-DR, Vα24, CXCR3, and CX3CR1 (**Figure 13**). Real-time compensation was performed at the instrument using FACSDiva V9.1, and data analysis and post-acquisition automatic traditional compensation were completed in FlowJo V10.10.

**Figure 13.**
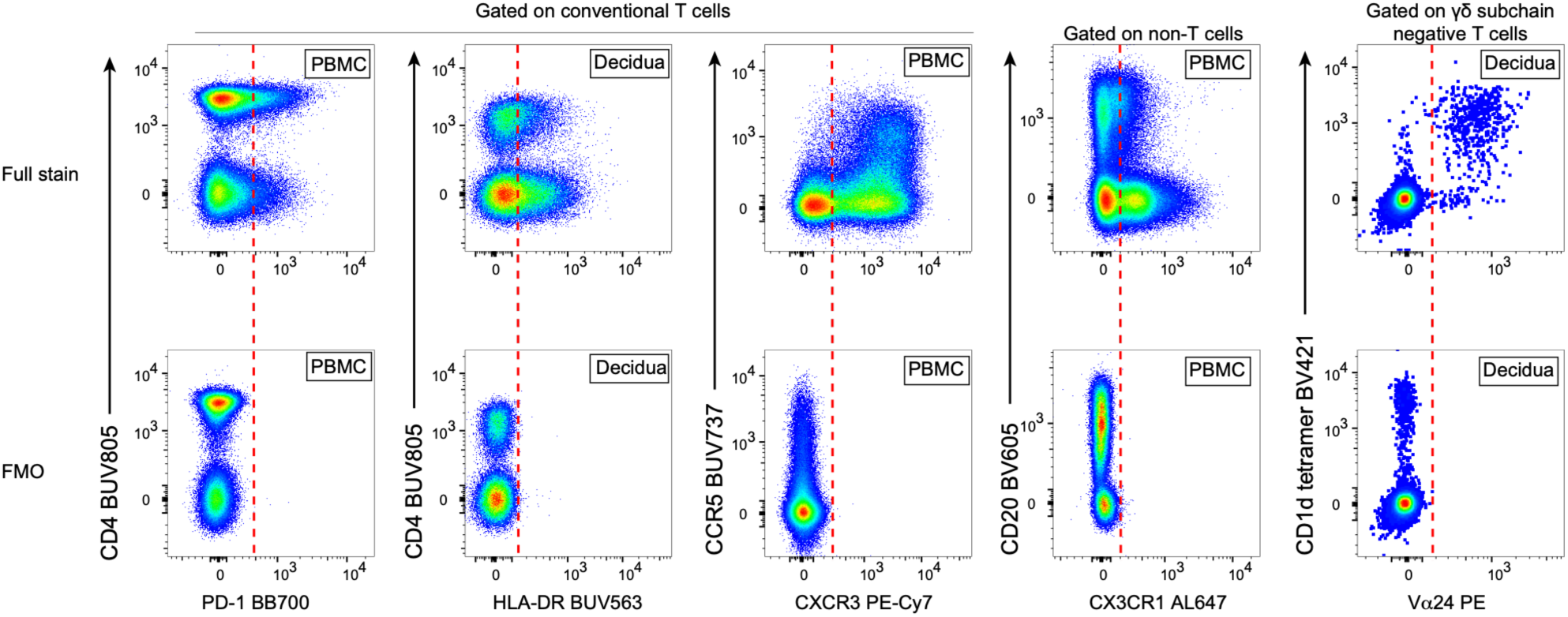
Fluorescence-minus-one (FMO) controls. FMO controls were used for PD-1, HLA-DR, CXCR3, CX3CR1, and Vα24. For each marker shown below, there are two representative dot plots. Plots in the bottom row are the FMO control sample, and plots in the top row are the fully stained samples. Red lines indicate the gate placement threshold, where gates were drawn slightly to the right of the negative population in the FMO and then applied to the full stain.

## Acknowledgements

The authors wish to thank Mario Roederer and Thomas Liechti for their troubleshooting support and advice during a puzzling experimental result. The Tulane National Biomedical Research Center (SCR_008167) Flow Cytometry Core’s (RRID: SCR_024611) staff Megan Varnado, Kaitlin Didier, Natalie Valencia provided excellent service and accommodated complicated experiments to complete this project during the COVID-19 pandemic. Lesli Sprehe and Elizabeth Scheef provided essential cell thawing expertise for critical experiments. We thank Diogo Magnani for maintaining and updating the Non-human Primate Reagent Resource (RRID:SCR_012986) website as a public database for investigators to share their reactivity findings.

## Data Availability Statement

The data that support the findings of this study are openly available in NIH ImmPort, Accession #SDY3667. The link to data is: https://immport.org/shared/study/SDY3367/summary.

## Ethical Statement

Rhesus macaque dams of Indian ancestry from the Tulane National Biomedical Research Center (TNBRC) were used for this study. All animal procedures were performed according to approved Institutional Animal Care and Use Committee protocols.

## References

1. Mostrom MJ, Scheef EA, Sprehe LM, Szeltner D, Tran D, Hennebold JD, et al. Immune Profile of the Normal Maternal-Fetal Interface in Rhesus Macaques and Its Alteration Following Zika Virus Infection. Front Immunol. 2021;12:719810.

2. Autissier P, Soulas C, Burdo TH, Williams KC. Immunophenotyping of lymphocyte, monocyte and dendritic cell subsets in normal rhesus macaques by 12-color flow cytometry: clarification on DC heterogeneity. J Immunol Methods. 2010;360(1-2):119–28.

3. Mahnke YD, Brodie TM, Sallusto F, Roederer M, Lugli E. The who’s who of T-cell differentiation: human memory T-cell subsets. Eur J Immunol. 2013;43(11):2797–809.

4. Huang Y, Zhu XY, Du MR, Li DJ. Human trophoblasts recruited T lymphocytes and monocytes into decidua by secretion of chemokine CXCL16 and interaction with CXCR6 in the first-trimester pregnancy. J Immunol. 2008;180(4):2367–75.

5. Tilburgs T, Roelen DL, van der Mast BJ, de Groot-Swings GM, Kleijburg C, Scherjon SA, et al. Evidence for a selective migration of fetus-specific CD4+CD25bright regulatory T cells from the peripheral blood to the decidua in human pregnancy. J Immunol. 2008;180(8):5737–45.

6. Vazquez J, Mohamed MA, Banerjee S, Keding LT, Koenig MR, Leyva Jaimes F, et al. Deciphering decidual leukocyte traffic with serial intravascular staining. Front Immunol. 2023;14:1332943.

7. DeJong CS, Maurice NJ, McCartney SA, Prlic M. Human Tissue-Resident Memory T Cells in the Maternal-Fetal Interface. Lost Soldiers or Special Forces? Cells. 2020;9(12).

8. Nogimori T, Moriishi E, Ikeda M, Takahama S, Yamamoto T. OMIP 075: A 22-color panel for the measurement of antigen-specific T-cell responses in human and nonhuman primates. Cytometry A. 2021;99(9):884–7.

9. Pitcher CJ, Hagen SI, Walker JM, Lum R, Mitchell BL, Maino VC, et al. Development and homeostasis of T cell memory in rhesus macaque. J Immunol. 2002;168(1):29–43.

10. DeGottardi MQ, Okoye AA, Vaidya M, Talla A, Konfe AL, Reyes MD, et al. Effect of Anti-IL-15 Administration on T Cell and NK Cell Homeostasis in Rhesus Macaques. J Immunol. 2016;197(4):1183–98.

11. Saravia J, Chapman NM, Chi H. Helper T cell differentiation. Cell Mol Immunol. 2019;16(7):634–43.

12. Zhu J. T Helper Cell Differentiation, Heterogeneity, and Plasticity. Cold Spring Harb Perspect Biol. 2018;10(10).

13. Mahnke YD, Beddall MH, Roederer M. OMIP-017: human CD4(+) helper T-cell subsets including follicular helper cells. Cytometry A. 2013;83(5):439–40.

14. Kara EE, Comerford I, Fenix KA, Bastow CR, Gregor CE, McKenzie DR, et al. Tailored immune responses: novel effector helper T cell subsets in protective immunity. PLoS Pathog. 2014;10(2):e1003905.

15. van de Berg PJ, Yong SL, Remmerswaal EB, van Lier RA, ten Berge IJ. Cytomegalovirus-induced effector T cells cause endothelial cell damage. Clin Vaccine Immunol. 2012;19(5):772–9.

16. Lee M, Lee Y, Song J, Lee J, Chang SY. Tissue-specific Role of CX(3)CR1 Expressing Immune Cells and Their Relationships with Human Disease. Immune Netw. 2018;18(1):e5.

17. Sakaguchi S, Mikami N, Wing JB, Tanaka A, Ichiyama K, Ohkura N. Regulatory T Cells and Human Disease. Annu Rev Immunol. 2020;38:541–66.

18. Munoz-Rojas AR, Mathis D. Tissue regulatory T cells: regulatory chameleons. Nat Rev Immunol. 2021;21(9):597–611.

19. Qian J, Zhang N, Lin J, Wang C, Pan X, Chen L, et al. Distinct pattern of Th17/Treg cells in pregnant women with a history of unexplained recurrent spontaneous abortion. Biosci Trends. 2018;12(2):157–67.

20. Tsuda S, Zhang X, Hamana H, Shima T, Ushijima A, Tsuda K, et al. Clonally Expanded Decidual Effector Regulatory T Cells Increase in Late Gestation of Normal Pregnancy, but Not in Preeclampsia, in Humans. Front Immunol. 2018;9:1934.

21. Svensson-Arvelund J, Mehta RB, Lindau R, Mirrasekhian E, Rodriguez-Martinez H, Berg G, et al. The human fetal placenta promotes tolerance against the semiallogeneic fetus by inducing regulatory T cells and homeostatic M2 macrophages. J Immunol. 2015;194(4):1534–44.

22. Salvany-Celades M, van der Zwan A, Benner M, Setrajcic-Dragos V, Bougleux Gomes HA, Iyer V, et al. Three Types of Functional Regulatory T Cells Control T Cell Responses at the Human Maternal-Fetal Interface. Cell Rep. 2019;27(9):2537–47 e5.

23. Robertson SA, Green ES, Care AS, Moldenhauer LM, Prins JR, Hull ML, et al. Therapeutic Potential of Regulatory T Cells in Preeclampsia-Opportunities and Challenges. Front Immunol. 2019;10:478.

24. Hartigan-O’Connor DJ, Poon C, Sinclair E, McCune JM. Human CD4+ regulatory T cells express lower levels of the IL-7 receptor alpha chain (CD127), allowing consistent identification and sorting of live cells. J Immunol Methods. 2007;319(1-2):41–52.

25. Godfrey DI, Uldrich AP, McCluskey J, Rossjohn J, Moody DB. The burgeoning family of unconventional T cells. Nat Immunol. 2015;16(11):1114–23.

26. Rossjohn J, Pellicci DG, Patel O, Gapin L, Godfrey DI. Recognition of CD1d-restricted antigens by natural killer T cells. Nat Rev Immunol. 2012;12(12):845–57.

27. Spada FM, Koezuka Y, Porcelli SA. CD1d-restricted recognition of synthetic glycolipid antigens by human natural killer T cells. J Exp Med. 1998;188(8):1529–34.

28. Guo J, Chowdhury RR, Mallajosyula V, Xie J, Dubey M, Liu Y, et al. gammadelta T cell antigen receptor polyspecificity enables T cell responses to a broad range of immune challenges. Proc Natl Acad Sci U S A. 2024;121(4):e2315592121.

29. Boyson JE, Rybalov B, Koopman LA, Exley M, Balk SP, Racke FK, et al. CD1d and invariant NKT cells at the human maternal-fetal interface. Proc Natl Acad Sci U S A. 2002;99(21):13741–6.

30. Terzieva A, Dimitrova V, Djerov L, Dimitrova P, Zapryanova S, Hristova I, et al. Early Pregnancy Human Decidua is Enriched with Activated, Fully Differentiated and Pro-Inflammatory Gamma/Delta T Cells with Diverse TCR Repertoires. Int J Mol Sci. 2019;20(3).

31. Rout N, Greene J, Yue S, O’Connor D, Johnson RP, Else JG, et al. Loss of effector and anti-inflammatory natural killer T lymphocyte function in pathogenic simian immunodeficiency virus infection. PLoS Pathog. 2012;8(9):e1002928.

32. Bond NG, Fahlberg MD, Yu S, Rout N, Tran D, Fitzpatrick-Schmidt T, et al. Immunomodulatory potential of in vivo natural killer T (NKT) activation by NKTT320 in Mauritian-origin cynomolgus macaques. iScience. 2022;25(3):103889.

33. Griffith JW, Sokol CL, Luster AD. Chemokines and chemokine receptors: positioning cells for host defense and immunity. Annu Rev Immunol. 2014;32:659–702.

34. Sallusto F, Mackay CR, Lanzavecchia A. The role of chemokine receptors in primary, effector, and memory immune responses. Annu Rev Immunol. 2000;18:593–620.

35. Nowacki TM, Kuerten S, Zhang W, Shive CL, Kreher CR, Boehm BO, et al. Granzyme B production distinguishes recently activated CD8(+) memory cells from resting memory cells. Cell Immunol. 2007;247(1):36–48.

36. Waterhouse NJ, Clarke CJ, Sedelies KA, Teng MW, Trapani JA. Cytotoxic lymphocytes; instigators of dramatic target cell death. Biochem Pharmacol. 2004;68(6):1033–40.

37. Tibbs E, Cao X. Emerging Canonical and Non-Canonical Roles of Granzyme B in Health and Disease. Cancers (Basel). 2022;14(6).

38. Jubel JM, Barbati ZR, Burger C, Wirtz DC, Schildberg FA. The Role of PD-1 in Acute and Chronic Infection. Front Immunol. 2020;11:487.

39. Wherry EJ, Kurachi M. Molecular and cellular insights into T cell exhaustion. Nat Rev Immunol. 2015;15(8):486–99.

40. Bajnok A, Ivanova M, Rigo J, Jr., Toldi G. The Distribution of Activation Markers and Selectins on Peripheral T Lymphocytes in Preeclampsia. Mediators Inflamm. 2017;2017:8045161.

41. Cibrian D, Sanchez-Madrid F. CD69: from activation marker to metabolic gatekeeper. Eur J Immunol. 2017;47(6):946–53.

42. Rea IM, McNerlan SE, Alexander HD. CD69, CD25, and HLA-DR activation antigen expression on CD3+ lymphocytes and relationship to serum TNF-alpha, IFN-gamma, and sIL-2R levels in aging. Exp Gerontol. 1999;34(1):79–93.

43. Shi H, Yang L, Zhang F, Zhou Y, Zhou Y. Diagnostic Value of CD25, CD69, and CD134 on Tuberculosis-Specific Antigen-Stimulated CD4+ T Cells for Tuberculous Pleurisy. J Immunol Res. 2023;2023:5309816.

44. Kumar BV, Ma W, Miron M, Granot T, Guyer RS, Carpenter DJ, et al. Human Tissue-Resident Memory T Cells Are Defined by Core Transcriptional and Functional Signatures in Lymphoid and Mucosal Sites. Cell Rep. 2017;20(12):2921–34.

45. Liechti T, Van Gassen S, Beddall M, Ballard R, Iftikhar Y, Du R, et al. A robust pipeline for high-content, high-throughput immunophenotyping reveals age- and genetics-dependent changes in blood leukocytes. Cell Rep Methods. 2023;3(10):100619.

46. Mahnke YD, Rajwa B. Gate Shape Matters. Cytometry Part A. 2026;109(3):165–6.

47. Bartmann C, Segerer SE, Rieger L, Kapp M, Sutterlin M, Kammerer U. Quantification of the predominant immune cell populations in decidua throughout human pregnancy. Am J Reprod Immunol. 2014;71(2):109–19.

48. Crespo ÂC, van der Zwan A, Ramalho-Santos J, Strominger JL, Tilburgs T. Cytotoxic potential of decidual NK cells and CD8+ T cells awakened by infections. J Reprod Immunol. 2017;119:85–90.

49. Tilburgs T, Claas FH, Scherjon SA. Elsevier Trophoblast Research Award Lecture: Unique properties of decidual T cells and their role in immune regulation during human pregnancy. Placenta. 2010;31 Suppl:S82-6.

50. Kieffer TEC, Laskewitz A, Scherjon SA, Faas MM, Prins JR. Memory T Cells in Pregnancy. Front Immunol. 2019;10:625.

51. Sacre K, Hunt PW, Hsue PY, Maidji E, Martin JN, Deeks SG, et al. A role for cytomegalovirus-specific CD4+CX3CR1+ T cells and cytomegalovirus-induced T-cell immunopathology in HIV-associated atherosclerosis. AIDS. 2012;26(7):805–14.

52. Berhanu D, Mortari F, De Rosa SC, Roederer M. Optimized lymphocyte isolation methods for analysis of chemokine receptor expression. J Immunol Methods. 2003;279(1-2):199–207.

53. Dolton G, Tungatt K, Lloyd A, Bianchi V, Theaker SM, Trimby A, et al. More tricks with tetramers: a practical guide to staining T cells with peptide-MHC multimers. Immunology. 2015;146(1):11–22.

54. Wooldridge L, Lissina A, Cole DK, van den Berg HA, Price DA, Sewell AK. Tricks with tetramers: how to get the most from multimeric peptide-MHC. Immunology. 2009;126(2):147–64.

55. Dambaeva SV, Durning M, Rozner AE, Golos TG. Immunophenotype and cytokine profiles of rhesus monkey CD56bright and CD56dim decidual natural killer cells. Biol Reprod. 2012;86(1):1–10.

